# A motor-based approach to induce chromosome-specific mis-segregations in human cells

**DOI:** 10.1101/2022.04.19.488790

**Authors:** My Anh Truong, Paula Cané-Gasull, Sippe G. de Vries, Wilco Nijenhuis, René Wardenaar, Lukas C. Kapitein, Floris Foijer, Susanne M.A. Lens

## Abstract

Various cancer types exhibit highly characteristic and recurrent aneuploidy patterns. The origin of these cancer type-specific karyotypes, and the extent to which they contribute to cancer progression, remains to be elucidated, partly because introducing or eliminating specific chromosomes in human cells still poses a challenge. Here, we describe a novel strategy to mis-segregate specific chromosomes at will in different human cell types. We employed Tet repressor (TetR) or nuclease dead Cas9 (dCas9) to link a plant-derived microtubule minus-end-directed kinesin (*Physcomitrella patens* Kinesin14VIb) to integrated Tet operon repeats and chromosome-specific endogenous repeats, respectively. By live- and fixed-cell imaging, we observed poleward movement of the targeted loci during (pro)metaphase. Kinesin14VIb-mediated pulling forces on the targeted chromosome were often counteracted by forces from kinetochore-attached microtubules. This tug of war resulted in chromosome-specific segregation errors during anaphase, and revealed that spindle forces can heavily stretch chromosomal arms. Using chromosome-specific FISH and single-cell whole genome sequencing, we established that motor-induced mis-segregations result in specific arm-level, and to a lesser extent, whole chromosome aneuploidies, after a single cell division. Our kinesin-based strategy to manipulate individual mitotic chromosomes opens up the possibility to investigate the immediate cellular responses to specific (arm level) aneuploidies in different cell types; an important step towards understanding how recurrent aneuploidy patterns arise in different cancer types.

## Introduction

Aneuploidy, defined as a chromosome number that is not the exact multiple of the usual haploid genome, is a prominent feature of cancer ^1^. Aneuploidy is the consequence of chromosomal instability (CIN), the increased frequency of chromosome segregation errors during mitosis that can result in gains or losses of entire chromosomes or of chromosomal arms in the daughter cells. Interestingly, different cancer types display different aneuploidy “signatures” with distinct whole-chromosome or arm level gains and losses. For instance, colorectal cancers frequently display gains of chromosome 7, 13 and 20q, and a loss of chromosome 18, whilst low grade gliomas are characterized by loss of 1p and gain of 19q ^2–7^. How these tissue-specific aneuploidy patterns arise is currently unclear, but potential mechanisms include non-random chromosome mis-segregation and selective pressures on cells with gains and losses of particular chromosomes. The relative contribution of each potential mechanism might differ between different tissues ^1^. Interestingly, conditions resulting in non-random chromosome mis-segregations have been identified in human cells ^8–10^. In addition, selective pressures that shape and stabilize the aneuploidy landscape over time have been described in yeast ^11–14^, as well as in human cells ^15–19^. However, how different cell types immediately respond and adapt to different chromosome gains and losses, and how specific aneuploidies drive or contribute to carcinogenesis, is still unknown. This is partly due to the technical challenge of introducing or eliminating distinctive chromosomes ^1^.

Various approaches have been developed to model specific whole-chromosome and arm-level aneuploidies, including microcell-mediated chromosome transfer ^20^, Cre-lox recombination of homologs to generate acentric and dicentric chromosomes ^21^, CRISPR/Cas9-mediated arm-level and whole-chromosome deletion ^7,22^, and centromere inactivation of chromosome Y by inducible degradation of CENP-A ^23^. Although these methods have generated valuable cell lines with particular chromosomal gains and losses ^24,25^, all of these approaches rely on clonal expansion, and therefore cells may have evolved during cell culture following the initial karyotype change. Complementary to these targeted studies, others have assessed the short-term consequences of random karyotype changes in human cells predominantly after mitotic checkpoint inhibition ^26–28^. We here introduce a novel approach to generate recent chromosome-specific aneuploidies in human cells, through targeted mis-segregation of specific chromosomes during mitosis. Leveraging the strong microtubule minus-end directed transport capacity of the spreading earthmoss (*Physcomitrella patens, Pp*) Kinesin14VIb ^29^, we managed to manipulate the orientation of a chromosome of interest on the mitotic spindle. Using TetR or nuclease-dead Cas9 (dCas9), we tethered *Pp* Kinesin14VIb to an integrated TetO repeat, or endogenous chromosome-specific repetitive loci, respectively. We show that the accumulation of this motor protein on either a subtelomeric repeat of chromosome 1p or a pericentromeric repeat of chromosome 9q, counteracts the congressional forces acting on the targeted sister-chromatids during early mitosis, and caused their specific mis-segregation during anaphase. Finally, we demonstrate that this strategy gives rise to daughter cells with specific arm-level gain or loss of the targeted chromosome(s) after one cell division cycle.

## Results

### Unequal distribution of a subtelomeric TetO locus in Chr1 through tethering of Kin14VIb

A pre-requisite for error-free chromosome segregation during mitosis is that all chromosomes align and bi-orient on the mitotic spindle ^30^. Bi-orientation and alignment are facilitated by the action of various plus-end directed microtubule-based motors that guide the movement of chromosomes towards the spindle equator ^30^. We hypothesized that by enriching minus-end-directed motors on a chromosome of interest, we could in turn transport that chromosome towards the spindle poles, causing it to misalign (Figure 1A, left). To test this hypothesis, we employed a minus-end directed kinesin from *P. patens*, Kinesin14VIb, which has several attractive properties. First, *Pp* Kinesin14VIb is the fastest known minus-end directed kinesin ^29^. Second, this plant motor shares very little homology with human kinesins ^31,32^, therefore *Pp* Kinesin14VIb overexpression is unlikely to interfere with their function. Third, we recently demonstrated that a truncated version of *Pp* Kinesin14VIb (amino acids 861-1321), lacking its cargo binding domain, could efficiently induce retrograde transport of organelles in human interphase cells ^31^. This truncated variant of *Pp* Kinesin14VIb was used in this study and is referred to as Kin14VIb.

**Figure 1:**
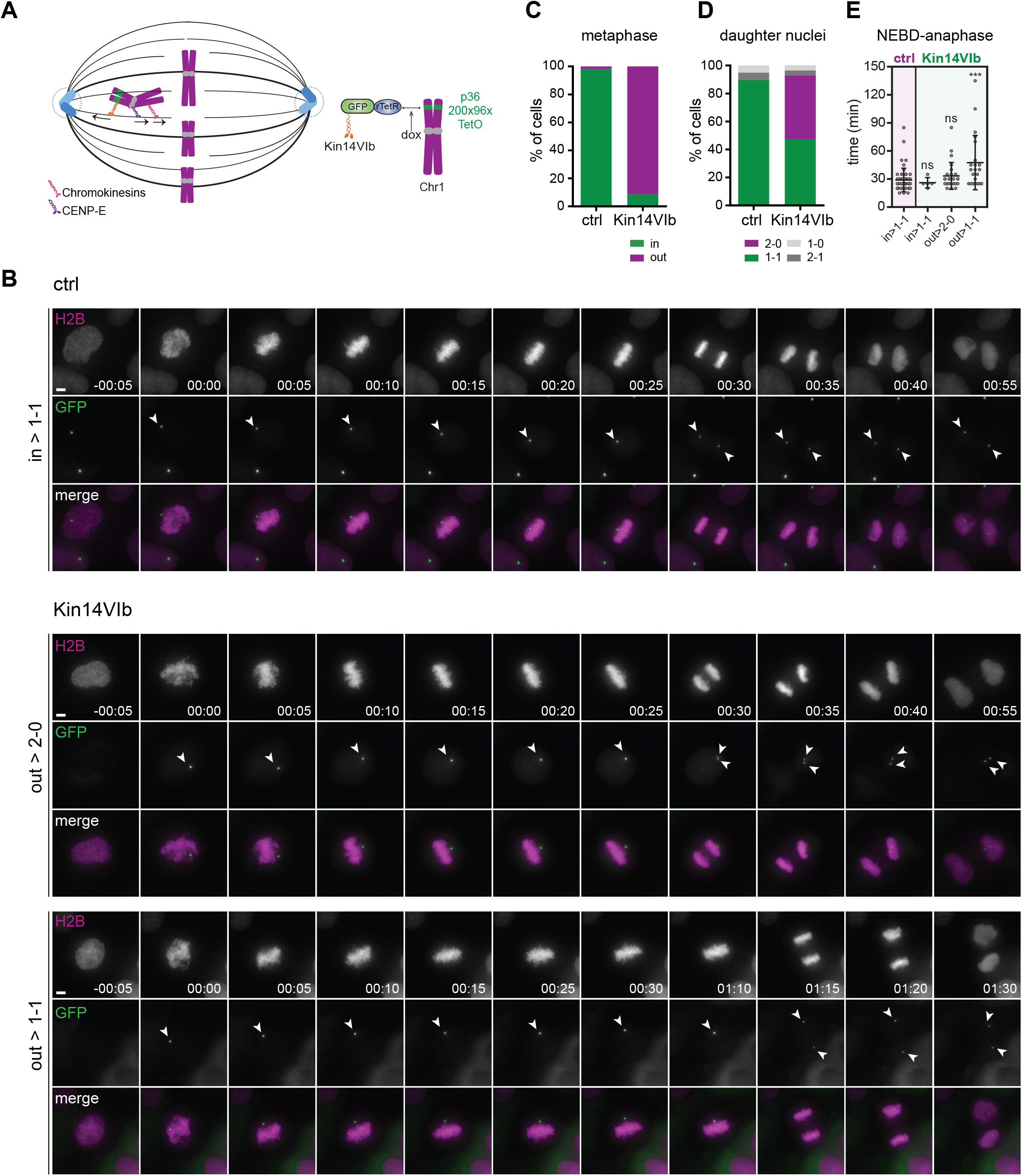
Consequences of Kin14VIb enrichment on a subtelomeric TetO repeat in Chr1. **A**. *Left*: Schematics of the rationale of the motor-based chromosome mis-segregation approach. Chromosome congression towards the spindle equator is facilitated by KT-MT attachments (not shown) and plus-end directed microtubule-based motors, such as chromokinesins at chromosome arms and CENP-E at kinetochores (KTs). The minus-end directed motor protein, Kin14VIb from *P. patens* (orange), is expected to induce poleward transport and counteract the plus-end directed motors. *Right:* In the U-2 OS-TetO cells, a 200×96× TetO repeat is integrated in a subtelomeric region of the p arm of one of the chromosomes 1^33^. Dimeric Kin14VIb is fused to GFP and rTetR, allowing doxycycline-inducible binding to the TetO repeat. **B-E**. Live cell microscopy of U-2 OS-TetO cells expressing rTetR-GFP (ctrl, n= 59) or rTetR-GFP-Kin14VIb (Kin14VIb, n=57)(Movies S1-3). **B**. Representative stills showing the most frequently observed metaphase localization and daughter cell distribution of the duplicated TetO locus (white arrowheads). Chromatin was visualized through expression of H2B-mCherry, GFP depicts the TetO locus. Time (h:min), scale bar = 5 µm. Due to the presence of a NES sequence in Kin14VIb, the TetO locus only becomes visible at NEB. **C**. Quantification of the metaphase localization of the TetO locus. Out = GFP focus observed outside the metaphase plate. **D**. Quantification of the distribution of the duplicated TetO locus over the daughter nuclei after anaphase. **E**. Time between nuclear envelop break down (NEB) and anaphase onset of the indicated categories of cells. Error bars depict mean ± SD. ns = not significant, *** p<0.005 (Mann-Whitney test).

To establish if Kin14VIb can promote minus-end-directed transport of a human mitotic chromosome, we made use of a U-2 OS cell line harboring a 200×96-mer TetO repeat in a subtelomeric region of one copy of Chr1p36 ^33^ (referred to as U-2 OS TetO, Figure 1A, right). To couple Kin14VIb to the chromosome with the TetO integration (referred to as the TetO chromosome), we fused the motor to the reverse Tet repressor and GFP (rTetR-GFP-Kin14VIb, Figure 1B). Expression of rTetR-GFP was used as control, and binding of rTetR-GFP or rTetR-GFP-Kin14VIb to the TetO locus was induced by doxycycline addition. By live cell imaging, we followed the trajectory of the TetO locus during mitosis (Figure 1B, movies S1-3). Because binding of rTetR-fusion proteins to TetO repeats can interfere with replication of this repeat during S phase ^34,35^, we added doxycycline immediately before filming to ensure that the captured mitotic cells were past S phase at the start of the experiment. In the majority of control cells, the TetO focus aligned on the metaphase plate and was subsequently separated during anaphase (Figure 1B, C). The two sister TetO foci segregated towards opposite spindle poles and eventually ended up in separate daughter nuclei (1-1 distribution, Figure 1B, D). In ∼90% of rTetR-GFP-Kin14VIb-expressing cells, however, the TetO focus was found outside the metaphase plate (Figure 1B, C). During the subsequent anaphase, the sister TetO foci co-segregated into one of the daughter nuclei, resulting in a 2-0 distribution of the TetO locus in ±45% of cell divisions (Figure 1 B, D). In addition, in ∼40% of dividing cells expressing rTetR-GFP-Kin14VIb, we observed that the TetO locus initially localized outside the metaphase plate, but eventually segregated equally into the two daughter nuclei in the subsequent anaphase (1-1 distribution, Figure 1B, bottom panel, C, D). Collectively, our live cell imaging data suggest that rTetR-GFP-Kin14VIb can mis-align a duplicated TetO locus during mitosis, and cause its subsequent unequal distribution across daughter nuclei.

### Kin14VIb-enrichment near telomeres does not prevent kinetochore-microtubule attachment

We observed a ∼30 min delay in anaphase onset in cells where the TetO locus initially resided outside the metaphase plate, but was segregated equally during anaphase (Figure 1E, category “out > 1-1”). However, the unequal (2-0) distribution of the duplicated TetO locus in Kin14VIb-expressing cells was not accompanied by a delay in anaphase onset (Figure 1E). This suggested that the kinetochores (KTs) of the Kin14VIb-bound TetO chromosome can acquire microtubule (MT) attachments that silence the mitotic checkpoint ^36,37^. To substantiate this, we performed immunofluorescence (IF) to assess TetO chromosome positioning and KT orientation in fixed U-2 OS cells synchronized in metaphase using the proteosome inhibitor MG132. Consistent with the live-cell imaging data, we detected the TetO locus inside the metaphase plate in nearly all control cells (Figure 2A, B). However, in ±80% of Kin14VIb-expressing cells, the TetO locus was found near one of the spindle poles, causing the p arms of the TetO sister chromatids to stick out from the metaphase plate (“arms out”, Figure 2A-C). Occasionally we observed the TetO locus to be stretched at the pole or to be disconnected from the chromosome (‘stretched/fragmented’, Figure 2B, C). Typically, we did not detect the centromere protein, CENP-C, in the vicinity of the TetO locus at the spindle poles. Instead, in ±76 % of cells displaying the TetO locus outside the metaphase plate, the KTs of the Kin14VIb-bound chromosomes were buried inside the metaphase plate together with the other aligned KTs (Figure 2C). In a few cases, we could distinguish the KTs of the TetO chromosome from the other KTs, with the inter-sister KT axis either orthogonal or near-parallel to the metaphase plate (Figure 2C). We then performed IF for the mitotic checkpoint protein Mad1, using its absence from KTs as a marker for MT attachment ^38^. When assessing the Mad1 status of KTs nearby the TetO locus, we did not find a difference between control cells (with TetO locus inside the metaphase plate) and Kin14VIb-expressing cells (with TetO locus outside the metaphase plate) (Figure 2D, S1). Moreover, in the few cells in which we could distinguish the KTs of the TetO chromosome, the sister KTs of this chromosome were predominantly attached (i.e. Mad1 negative), irrespective whether the inter-sister KT axis was orthogonal or near-parallel to the metaphase plate (Figure 2D, S1).

**Figure 2:**
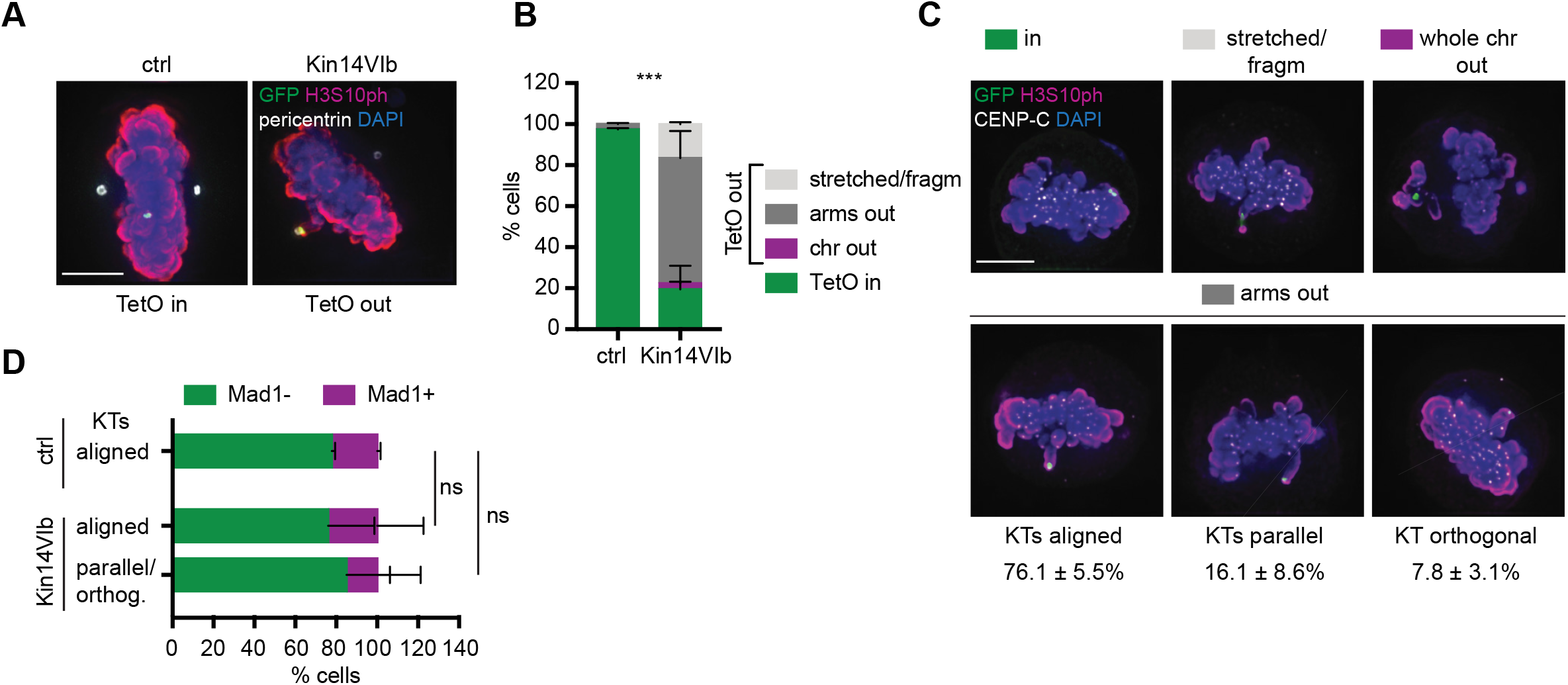
Metaphase orientation and KT attachment status of the Kin14VIb-bound TetO chromosome. **A**. IF for pericentrin, GFP, H3S10ph of U-2 OS TetO cells in metaphase. The TetO locus (GFP focus) is aligned on the metaphase plate (TetO in) in control condition (rTetR-GFP), and co-localizing with one of the two pericentrin-marked centrosomes (TetO out) in rTetR-GFP-Kin14VIb-expressing cells. **B, C**. IF for CENP-C, GFP, H3S10ph of U-2 OS TetO cells in metaphase and expressing rTetR-GFP-Kin14VIb. **B**. Quantification of the fraction of cells with the indicated orientation of the duplicated TetO locus or TetO chromosome. Experiment was performed in duplicate (mean ± SD, n ≥ 29 per condition, per experiment). *** p<0.001 (Fisher’s exact test, comparing the fraction of cells with the TetO locus in vs. out in ctrl vs. Kin14VIb expressing cells). **C**. Representative images of the different orientations of the TetO chromosome and the sister KTs are shown. DNA was visualized by DAPI. Scale bar = 5 µm. Numbers below the images depict the frequency of the indicated orientation of the sister-KTs of the duplicated TetO chromosome with its p arm sticking out towards the spindle pole (arms out). KT orthogonal: inter-sister KT axis (dotted line) perpendicular to metaphase plate; KT parallel: inter-sister KT axis is (near-)parallel the metaphase plate; KT aligned: sister KTs buried inside the metaphase plate. **D**. Quantification of Mad1 status of KTs nearby the TetO locus. Representative IF images are shown in Figure S1. In case the KTs of the TetO chromosome were aligned, we scored whether KTs in the vicinity of the TetO locus were Mad1+. In case the KTs of the TetO chromosome could be distinguished in rTetR-GFP-Kin14VIb-expressing cells (with either orthogonal or parallel oriented KTs), we scored if at least one of the KTs was Mad1+. Experiments were performed in duplicate (mean ± SD, n ≥ 27 per condition, per experiment). Ns = not significant (χ^2^ test, comparing the fraction of cells with the Mad1-vs. Mad1+ in ctrl cells vs. Kin14VIb cells with aligned KTs, or ctrl cells vs. Kin14VIb cells with orthogonal/parallel KTs).

Thus, recruiting minus-end directed kinesins near chromosomal telomeres did not prevent KTs from binding microtubules. Notably, Kin14VIb pulled the p arms of the sister chromatids, but not the entire duplicated chromosome, towards one of the spindle poles. This implies that the sister KTs were not syntelically attached (i.e. both sister-KTs attached by MTs coming from the same spindle pole, Figure S1B) to MTs derived from the spindle pole to which the TetO locus was transported, as this would have caused the entire chromosome to remain near that spindle pole. It is therefore more likely that the sister-KTs of the TetO chromosome were either attached in an amphitelic (sister-KTs attached by MTs deriving from opposite spindle poles) or merotelic fashion (one of the sister-KTs attached by MTs coming from opposite spindle poles, Figure S1B), or were potentially syntelically attached by MTs from the opposite spindle pole.

### On-target mis-segregation of the Kin14VIb-bound TetO chromosome during anaphase

To understand how simultaneous pulling by KT-attached MTs and Kin14VIb would affect chromosome orientation and segregation during anaphase, we analyzed fixed anaphases of control cells with a 1-1, or of Kin14VIb-expressing cells with a 2-0 distribution of the TetO locus. We observed that ∼35% of rTetR-GFP expressing cells displayed chromosome bridges or lagging chromosomes, illustrating the rate of ongoing CIN in U-2 OS cells, as previously reported ^39^. However, in the rTetR-GFP-Kin14VIb-expressing cells with a 2-0 distribution of the TetO locus, we observed a chromosome bridge or lagging chromosome in >90% of the cells, indicating that tethering of the kinesin to the TetO chromosome significantly increased CIN. Interestingly, in contrast to control cells, the bridges and laggards in Kin14VIb-expressing cells frequently included at least one lagging chromosome or a stretch of chromatin positioned along the center of the spindle axis (mid-axis, white dashed line, Figure 3A). Moreover, in ± 50% of Kin14VIb-expressing cells with a 2-0 distribution of TetO sister foci, we observed a stretched chromatid arm, with the TetO locus residing near one of the spindle poles and its KT lagging behind the two main masses of segregating chromosomes (Figure 3A-C). Occasionally, we observed the arms of both sister chromatids being stretched in Kin14VIb-expressing cells, most likely reflecting syntelic attachment of the sister KTs by MTs coming from the spindle pole, opposite from the one to which the TetO locus is transported (Figure 3B, Figure S2A, B). Of note, these typical stretched arms were never observed in control cells, and resided along the spindle mid-axis (Figure 3A, B, Figure S2C), presumably because the Kin14VIb-bound TetO focus was transported towards the spindle pole. Thus, Kin14VIb-mediated poleward transport of a subtelomeric TetO locus induces a specific type of segregation error in anaphase: a heavily stretched p arm of a single sister chromatid. Our data suggest that while each p arm of the duplicated TetO chromosome is pulled toward one spindle pole by Kin14VIb, one KT of this duplicated chromosome is attached to MTs coming from the opposite spindle pole. This leads to a single sister chromatid arm stretching and lagging during anaphase (Figure 3C, S2C).

**Figure 3:**
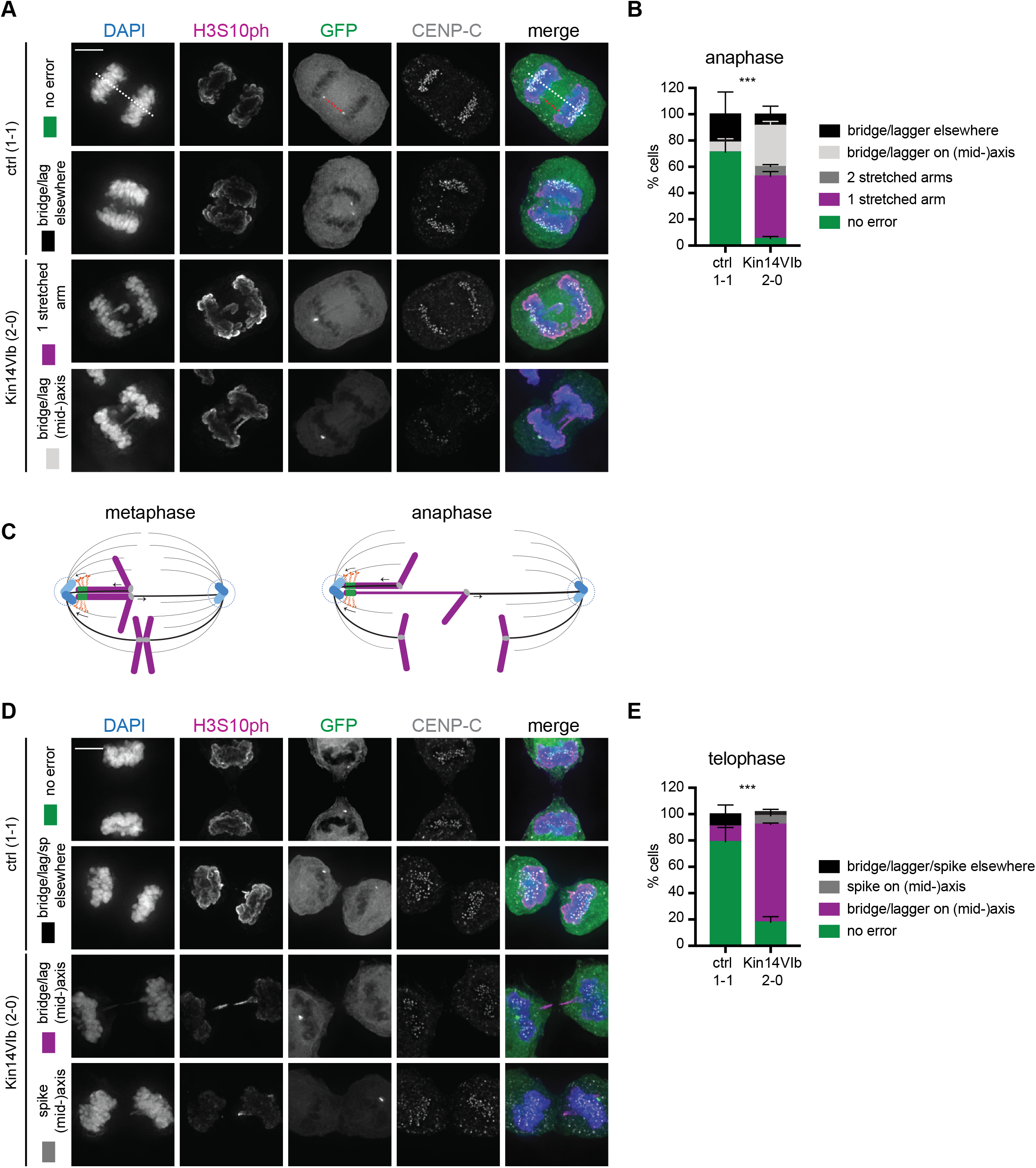
Opposing pulling forces acting on the Kin14IVb-bound TetO chromosome causes its stretching and unequal segregation during anaphase. **A, D**. IF for CENP-C, GFP, H3S10ph of U-2 OS TetO cells in anaphase (A) and telophase (B) expressing either rTetR-GFP (ctrl) or rTetR-GFP-Kin14VIb. Representative images of the most frequently observed anaphase and telophase errors in cells with either a 1-1 (majority of events in ctrl), or 2-0 (majority of events in Kin14VIb) TetO locus distribution. DNA was visualized with DAPI. Scale bar = 5 µm. White dashed line depicts the pole-pole mid-axis, while the red dashed line indicates the TetO-TetO axis (A, 1^st^ row). **B, E**. Frequencies of the different types of segregation errors observed during anaphase (B) and telophase (E). To define anaphase and telophase segregation errors involving the TetO chromosome, errors along the TetO-TetO axis or pole-pole mid-axis (collectively called: (mid-)axis), were distinguished from errors that were observed to occur not along one of these axis (bridge/lagger elsewhere). Experiments were performed in duplicate (mean ± SD, n ≥ 23 per condition, per experiment). *** p<0.001 (Fisher’s exact test, comparing the fraction of cells displaying no error vs. 1 or 2 arms stretched in ctrl vs. Kin14VIb cells in anaphase (B), or cells displaying no error vs. bridge/lagger/spike on the (mid-)axis in ctrl vs. Kin14VIb cells in telophase (E)). **C**. Schematic representation of the most frequently observed consequence of Kin14VIb binding to the subtelomeric TetO locus of chromosome 1 in U-2 OS-TetO cells during metaphase and anaphase.

### Tethering of Kin14VIb to a subtelomeric TetO repeat in Chr1 results in 1p copy number alterations after a single cell division

Depending on the counteracting forces exerted by the KT-attached MTs and the TetO-bound Kin14VIb, we anticipated that the stretched sister chromatid would either be mis-segregated towards the TetO locus, or be further stretched by spindle forces, with the KT moving towards one spindle pole, and the TetO locus towards the opposite spindle pole, and causing the p arm to cross and persist in the cytokinetic furrow. Analysis of telophases with a 1-1 and 2-0 distribution of the TetO locus revealed that the 2-0 distribution was frequently (75%) accompanied by a histone H3 Serine10 phosphorylated (H3S10ph+) chromatin bridge inside the furrow, or by a spike of H3S10ph+ chromatin in at least one of the daughter nuclei (Figure 3D, E, Figure S2C). H3S10 is a substrate of the Aurora B kinase, and phosphorylation of this histone site during telophase suggests close proximity to, or recent passing through, the spindle midzone and midbody, structures on which Aurora B localizes during this phase of the cell cycle ^40,41^(Figure S2C). After FISH probe labelling for Chr1 centromeres (Chr1-CEN), we assessed how frequently the Chr1 centromere would be unequally segregated in Kin14VIb-expressing cells. We observed an unequal distribution of Chr1 centromeres in ±35% of the Kin14VIb expressing cells (Figure 4A, B). Note that U-2 OS is triploid for Chr1 (Figure 4A, S3A), and that we did not select for cells with a 2-0 TetO distribution since we were unable to reliably combine IF with FISH. The unequal Chr1-CEN distribution observed in Kin14VIb-expressing cells included cases where Chr1-CEN was unequally distributed over the two daughter nuclei, but more frequently included anaphases with Chr1-CEN foci appearing as lagging or fragmented (Figure 4A, B). The combined IF and centromere FISH data suggest that Kin14VIb binding close to the telomere may cause a whole Chr1 mis-segregation and subsequent aneuploidy in ±11% of cell divisions, and most likely 1p alterations after the vast majority of cell divisions. Accordingly, shallow whole genome sequencing of single G1 nuclei (scKaryo-Seq) of U-2 OS TetO cells, revealed an increased mis-segregation rate for Chr1p specifically in rTetR-GFP-Kin14VIb transduced cells, evidenced by an increased aneuploidy score for this chromosome arm (Figure 4C, S3). Note that all chromosome arms displayed variations due to the overall heterogenous nature of U-2 OS karyotypes, but that 1p exhibited the highest increase in aneuploidy score amongst all chromosome arms when rTetR-GFP-Kin14VIb was transduced (Figure 4C, Supplemental Table 1). Although the Chr1-CEN FISH and single cell WGS data suggest that the Chr1p arm breaks during cell division, we did not detect an increase in the double strand break (DSB) marker γH2AX on the TetO locus, nor on the H3S10ph+ chromatin bridges in Kin14VIb-expressing telophase cells (Figure S4) ^42^. This suggests that persistent stretching of the p arm per se does not cause detectable DNA damage, and that the actual breakage occurs later (e.g. during abscission)^43,44^, or that the DNA damage is detected later (e.g. in G1)^45^. To conclude, our data suggest that Kin14VIb-mediated poleward transport of a subtelomeric repeat in Chr1p causes stretching of a single chromatid arm and subsequent breakage of this arm after mitosis, leading to copy number alterations of Chr1p in the resulting daughter cells.

**Figure 4:**
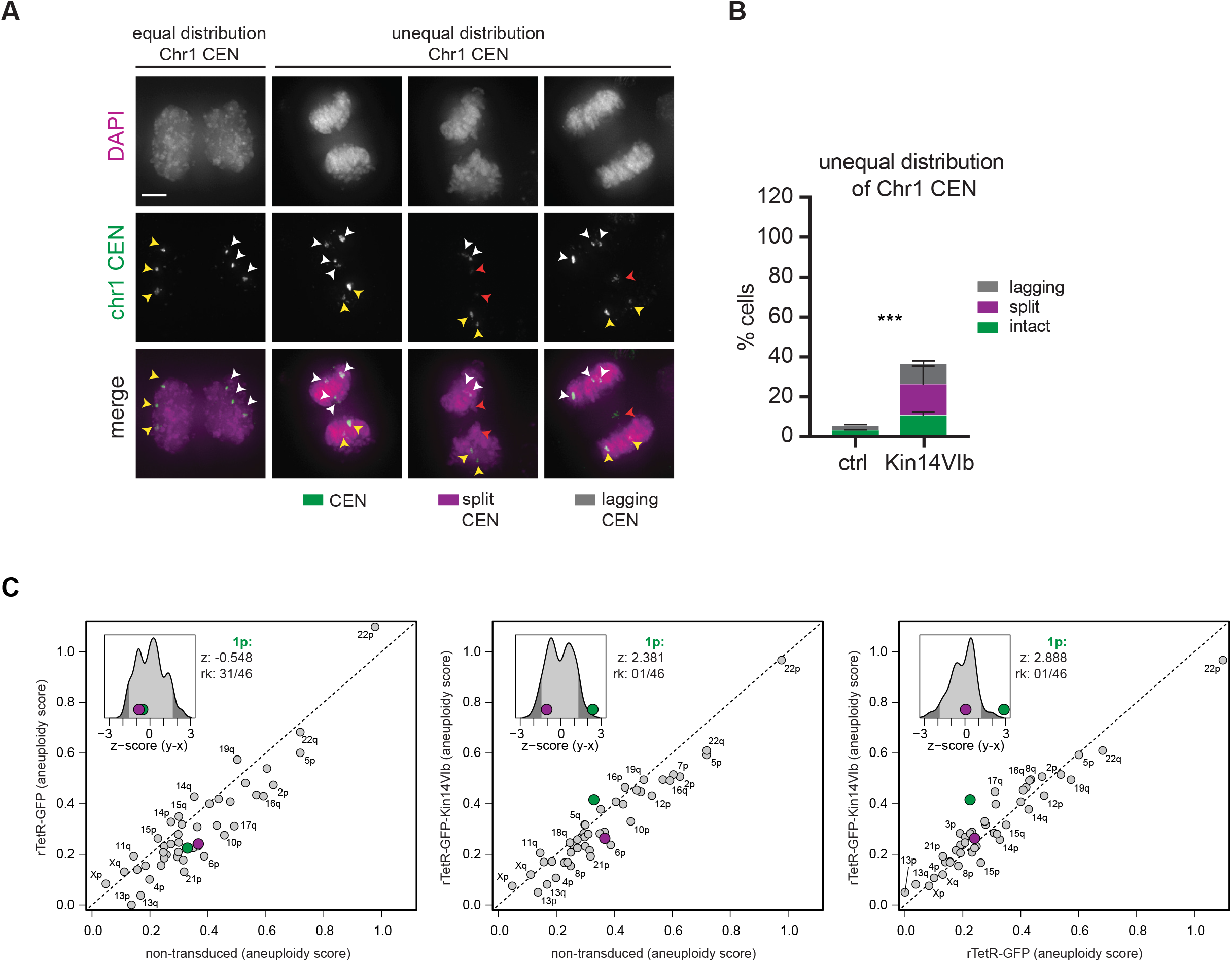
Tethering of Kin14VIb to a subtelomeric TetO repeat in Chr1p leads to 1p copy number alterations after a single cell division. **A**. Representative images of U-2 OS TetO anaphases (rTetR-GFP, ctrl or rTetR-GFP-Kin14VIb expressing) with FISH-marked centromeres of chromosome 1 (chr1 CEN). White and yellow arrowheads: chr1 CEN in 2 daughter nuclei; red arrowheads: chr1 CEN fragmented or lagging between the two daughter nuclei. **B**. Frequency of the different types of errors observed in cases of an “unequal distribution” in A. Experiment was performed in duplicate (mean ±SD, n ≥ 45 per condition, per experiment). *** p<0.001 (Fisher’s exact test comparing the fraction of cells displaying an equal vs. unequal distribution of chr1 CEN FISH foci in ctrl vs. Kin14VIb cells). **C**. Plots comparing aneuploidy scores of all chromosomal p and q arms between non-transduced U-2 OS TetO cells and rTetR-GFP-transduced cells (left), non-transduced cells and rTetR-GFP-Kin14VIb-transduced cells (middle), and rTetR-GFP-transduced and rTetR-GFP-Kin14VIb-transduced cells (right). Aneuploidy scores were calculated from the single cell WGS data of single G1 nuclei following a round of mitosis with doxycyclin-induced rTetR-TetO binding (see materials and methods). The median copy number of each bin (median across libraries) was determined for the non-transduced U-2 OS TetO condition and used as reference (see Figure S3A). The aneuploidy scores of the chromosomal arms are deviations from this reference (average absolute difference of the bins that are associated with the arm). Inserts show density plots of the difference of the scores of the two conditions that are compared (y-axis minus x-axis). These differences are expressed as z-scores (number of standard deviations from the mean). The 95% confidence interval is depicted in light grey. The values of the 1p and 1q arm are indicated with a green and purple dot respectively. Z-score value for 1p and its ranking (rk) is indicated (see Supplemental Table 1).

### Harnessing dCas9 to link Kin14VIb to endogenous chromosome-specific DNA repeats in RPE1 cells

Having established a tool to robustly induce the specific mis-segregation of single chromatid arms, we aimed to make this technology more broadly applicable, by removing the dependence on chromosomal integration of exogenous repetitive DNA sequences, thereby facilitating its introduction into more physiological model systems. To this end, we developed a strategy to link Kin14IVb to chromosome-specific endogenous repeats using dCas9 ^46^. To tether Kin14VIb to dCas9 on a targeted endogenous locus, we made use of FK506 binding protein 12 (FKBP12) and FKBP-rapamycin binding (FRB) domain, which can be chemically induced to dimerize upon addition of rapalog ^47,48^(Figure 5A). Additionally, to increase Kin14VIb-mediated pulling efficiency, we attached the dimerization domain GCN4 to fluorescent FRB-Kin14VIb, thereby clustering Kin14VIb into dimer of dimers, and rendering the motor highly processive, independently of cargo binding ^29,31^. We then generated RPE1 (non-transformed, near-diploid) cell lines stably expressing low levels of dCas9-GFP-3xFKBP and high levels of FRB-mCherry-GCN4-Kin14VIb, the latter controlled by a doxycyclin-inducible promotor (see material and methods; this cell line is referred to as RPE1 dCas9-Kin14VIb). In agreement with earlier observations^31^, GCN4-Kin14VIb weakly decorated the spindle poles, occasionally resulting in dCas9-GFP-3xFKBP localization at these sites (movies S4-8).

**Figure 5:**
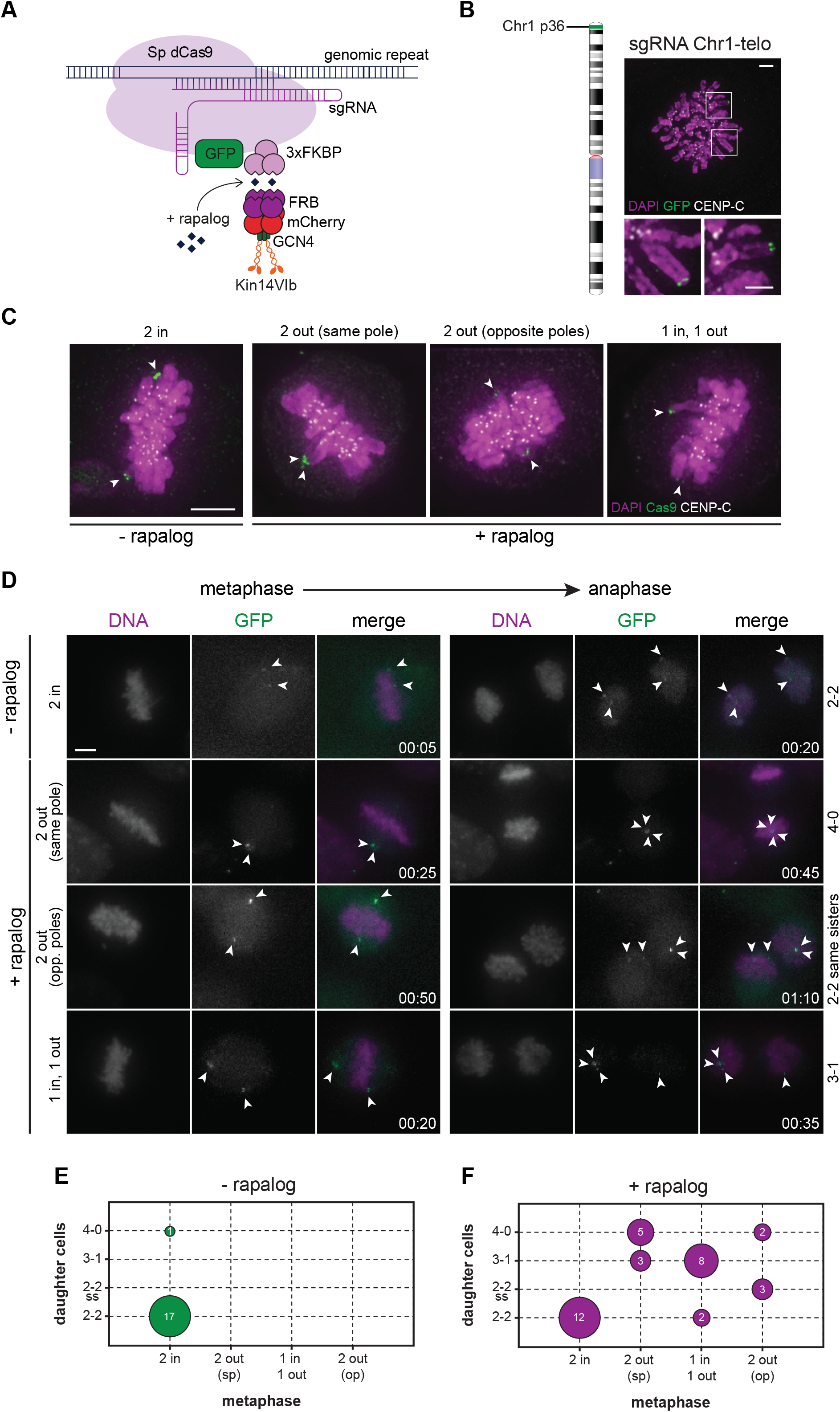
Tethering Kin14VIb to endogenous subtelomeric DNA repeats of Chr1p in RPE1 cells. **A**. Schematic representation of the strategy to couple FRB-mCherry-GCN4-Kin14VIb to endogenous chromosome-specific loci in a dCas9-GFP-3xFKBP expressing RPE1 cell line. **B**. The Chr1-telo sgRNA binds to a subtelomeric repetitive DNA sequence in the p-arm of chromosome 1 (left). Chromosome spread of RPE1 cells expressing dCas9-GFP-3xFKBP and transduced with Chr1-telo sgRNAs (right). Magnifications of the white boxed regions (each region showing one Chr1 homolog) are shown below the IF image (scale bar = 2 µm). **C**. IF for Cas9 and CENP-C of metaphases of RPE1-dCas9-Kin14VIb cells transduced with Chr1-telo sgRNA in the presence or absence of 500 nM rapalog. The Chr1-telo loci (Cas9 foci) are aligned on the metaphase plate in control condition (-rapalog), whilst the loci are facing either a single or opposite poles after rapalog addition to induce Kin14VIb binding to dCas9. **D,E**. Live cell microscopy of asynchronously growing RPE1-dCas9-Kin14VIb cells transduced with Chr1-telo sgRNA in the presence or absence of 500 nM rapalog to induce kinesin binding to the subtelomeric locus. **D**. Stills showing the most frequently observed metaphase localization and daughter cell distribution (white arrowheads) of the duplicated Chr1-telo loci. SiR-DNA was used to visualise the DNA, GFP depicts the Chr1-telo loci. Time (h:min). Scale bar = 5 µm. Note, that in the upper panel, the cell was already in mitosis when imaging started (t =0:00). **E**. Plots showing the relationship between the indicated metaphase orientation of the Chr1-telo loci, and the indicated distributions of the loci in the daughter cells. Circle size reflects relative cell numbers. The actual cell numbers per condition are indicated in the circles. Cells were derived from two independent imaging experiments. N = 18 cells (-rapa), n = 35 cells (+ rapa). Sp = same pole, op = opposite pole, ss = same sisters.

To test if kinesin enrichment on a endogenous subtelomeric repeat would have similar consequences for the targeted chromosome as motor binding to a large integrated exogenous subtelomeric repeat (as in the U-2 OS TetO cells), we transduced RPE1 dCas9-Kin14VIb cells with a sgRNA targeting dCas9 to a subtelomeric DNA repeat in Chr1p36 (Chr1-telo, Figure 5B)^8,49^. After rapalog addition, metaphase-arrested RPE1 cells displayed the typical phenotype of chromosomal arms sticking out from the metaphase plate with the dCas9-bound subtelomeric locus facing the spindle pole (Figure 5C), similar as observed in U-2 OS TetO cells expressing Kin14VIb. We next followed the alignment and segregation of the two Chr1-telo loci over time by tracking the GFP foci throughout mitosis, by live cell imaging immediately after addition of rapalog (Figure 5D). While the majority of control cells showed complete alignment of both Chr1-telo loci in metaphase, rapalog-treated cells frequently displayed at least one Chr1-telo locus outside the metaphase plate (Figure 5D, E). Similar to what we observed in fixed cells (Figure 5C), three distinct categories of foci residing “outside” the metaphase plate could be distinguished: 1) one of the foci aligned while the other was located near a spindle pole, 2) both foci resided near the same pole, or 3) the two foci were found near opposite spindle poles (Figure 5D, E). In the subsequent anaphase, these orientations mostly led to a Chr1-telo locus distribution between daughter nuclei of respectively 3-1, 4-0, or 2-2 with sister loci ending up in the same daughter nuclei (2-2, same sisters) (Figure 5E). Although efficiency varied per experiment, an unequal distribution of the Chr1-telo loci was consistently observed in the rapalog-treated RPE1 cells (Figure 5E).

### Inducing targeted CIN of Chr9q in RPE1 cells

The versatile nature of the dCas9-based system in RPE1 cells allowed targeting of Kin14VIb to other chromosomes and to other chromosomal regions. We therefore next studied the consequences of Kin14VIb binding to a chromosomal region more proximal to the kinetochore, by transducing a sgRNA specific to a pericentromeric DNA repeat of Chr9q (Chr9-cen, Figure 6A, B)^50,51^. Since this repeat is predicted to harbor ∼550,000 Chr9-cen sgRNA binding sites, while the subtelomeric repeat in Chr1p is predicted to harbor ∼1,400 Chr1-telo sgRNA binding sites (Tovini et al., accompanying manuscript), Chr9-cen GFP-foci appear larger than the Chr1-telo GFP-foci (Figure 5B vs 6B). Live-cell microscopy showed that both Chr9-cen homologs were most frequently transported towards either the same or opposite spindle pole (Figure 6C, D). Consequently, this recurrently led to daughter cells with a 4-0 or a 2-2 (same sisters) distribution of Chr9-cen foci (Figure 6D). Note that in the absence of rapalog, ±30% of cells already displayed some Chr9-cen specific anaphase errors (lagging or bridging/fragmented Chr9-cen foci) sometimes leading to 3-1 focus distribution in the daughter cells (Figure 6C, D). This might be a consequence of incomplete replication of the repeat caused by binding of dCas9 to the pericentromeric repeats on Chr9 during S phase^52^. Despite this higher background anaphase error rate, coupling Kin14Vib to Chr9-cen repeats clearly induced an increase in the unequal distribution of the Chr9-cen locus during mitosis.

**Figure 6:**
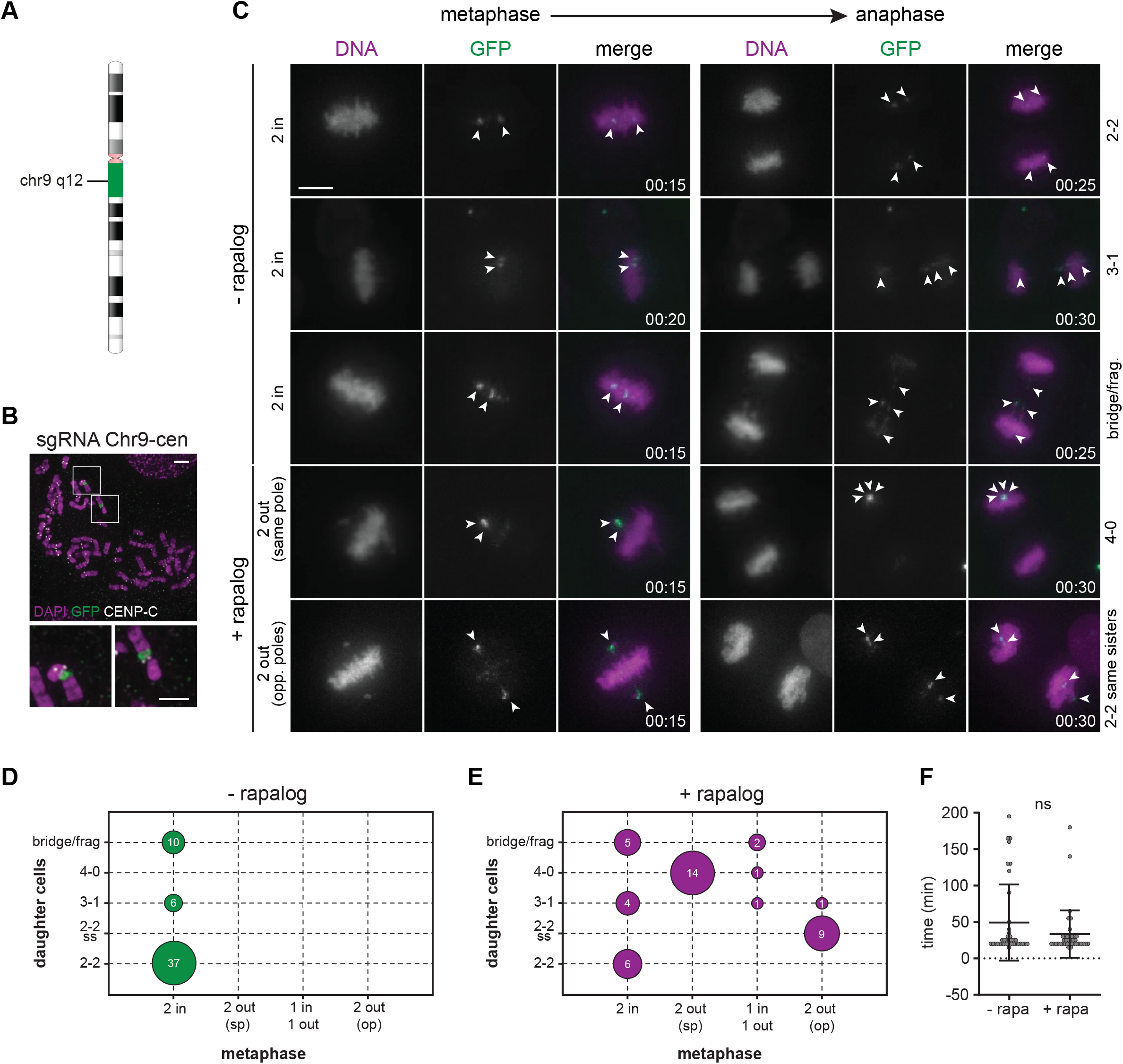
Coupling Kin14IVb to endogenous pericentromeric repeats of Chr9q in RPE1 cells. **A**. The Chr9-cen sgRNA binds to a pericentromeric repetitive DNA sequence in the q arm of chromosome 9 (upper). **B**. Chromosome spread of RPE1 cells expressing dCas9-GFP-3xFKBP and transduced with Chr9-cen sgRNAs. Magnifications of the white boxed regions (each region showing one mitotic Chr9 homolog) are shown below the IF image (scale bar = 2 µm). **C-G**. Live cell microscopy of asynchronously growing RPE1-dCas9-Kin14VIb cells transduced with Chr9-cen sgRNA and plus/minus rapalog to induce kinesin binding to the pericentromeric locus (Movies S4-S8). **C**. Representative stills showing the most frequently observed metaphase localization and daughter cell distribution (white arrowheads) of the duplicated Chr9-cen loci. SiR-DNA was used to visualise the DNA, GFP depicts the Chr9-cen loci. Time (h:min). Scale bar = 5 µm. **D**. Frequency of the observed metaphase localizations of the Chr9-cen loci as shown in (C). *** p<0.001 (Fisher’s exact test, comparing the fraction of cells displaying 2 Chr9-cen loci inside vs. at least 1 Chr9-cen locus outside the metaphase plate in the absence or presence of rapalog). **E**. Subsequent distribution of the duplicated Chr9-cen loci over the daughter nuclei during anaphase. Experiment was performed in duplicate (mean ± SD, n ≥ 15 cells per experiment -rapa, and n ≥ 13 per experiment +rapa). *** p<0.001 (Fisher’s exact test, comparing the fraction of cells with equal (2-2) vs. unequal (3-1, 4-0, or 2-2, same sisters) distribution of Chr9-cen loci in the absence or presence of rapalog). (**D**) Plots showing the relationship between the indicated metaphase orientation of the Chr1-telo loci, and the indicated distributions of the loci in the daughter cells. Circle size reflects relative cell numbers. The actual cell numbers per condition are indicated in the circles. Cells were derived from two independent imaging experiments. N = 53 cells (-rapa), n = 43 cells (+ rapa). Sp = same pole, op = opposite pole, ss = same sisters. **E**. Time between nuclear envelop break down (NEB) and anaphase onset in the presence (Kin14VIb bound to dCas9) or absence (Kin14VIb expressed, but not bound to dCas9) of rapalog. Ns = not significant (Mann-Whitney test). Dots represent individual cells obtained from two independent imaging experiments (n = 38 cells for both conditions).

### Tethering Kin14VIb to a pericentromeric DNA repeat in Chr9 results in 9q aneuploidies after a single cell division

Detailed IF analysis of fixed mitoses revealed some striking differences in the consequences of Kin14VIb binding near telomeres (Figure 2A-C, 5C) or near centromeres (Figure 7). In Chr9-cen sgRNA expressing cells (plus rapalog), we frequently observed at least two entire sister chromatid arms near the spindle pole(s) that lacked CENP-C (Figure 7A-C), suggesting that only the q arms on which dCas9 and Kin14VIb were bound, were transported towards the spindle pole(s). Moreover, instead of chromosomal arms sticking out from the metaphase plate, as observed when Kin14VIb is bound to the Chr1-telo loci (Figure 2A-C, 5C), we found that the sister chromatid 9q arms at the poles were connected to chromosomes in the metaphase plate by a stretch of dCas9-bound chromatin (Figure 7A). Our data suggest that while the 9q arms were transported towards the spindle poles by Kin14VIb attached to the pericentromeric region, the sister KTs of this chromosome congressed towards the metaphase plate, most likely via microtubule interactions. This could explain the absence of a mitotic delay despite visible poleward transport of Chr9-cen loci (Figure 6E). Since we observed stretching of Chr9-cen chromatin as early as metaphase, we predict that this stretch of pericentromeric heterochromatin may persist as a (fine) chromatin bridge during anaphase and eventually break during telophase or later, causing an unequal distribution of the q arm over the daughter cells (Figure 7D-F, 8A). To reveal the fate of Chr9 after a round of Kin14VIb-induced mis-segregation, we single cell sorted EdU+ RPE1-dCas9-Kin14VIb G1 cells following an overnight incubation with rapalog and processed the cells for scKaryo-seq. As expected, the single cell karyotype heatmaps of the near-diploid CIN-RPE1 cells were more homogeneous than those derived from CIN+ U-2 OS cells (Figure 8B, Figure S3). In addition to Chr10q, a clonal abnormality specific to RPE1 cells^9,28^, RPE1-dCas9-Kin14VIb cells displayed a clonal gain of Xq that arose during clonal selection required to derive the dCas9-Kin14VIb expressing cell line (Figure 8B). In addition to this, we observed an infrequent gain of 19p, which somehow increased after sgRNA transduction and Kin14VIb expression. Importantly, the rapalog-inducible FKBP12-FRB dimerization modality of our system allowed us to directly assess the effect of Kin14VIb motor binding to Chr9, on Chr9 distribution after cell division. Indeed, we detected Chr9 copy number deviations in ∼24% (25/105) of the cells treated with rapalog versus ∼10% (4/39) of the cells that were not treated with rapolog (Figure 8B). Note that the “2-2, same sisters” type of mis-segregation that we observed after microscopic inspection (Figure 6D, 7), is unlikely to result in a copy number deviation, and will thus be missed by scKaryo-seq. Interestingly, the addition of rapalog increased the aneuploidy score of 9q (Figure 8C, Supplemental Table 2), and resulted in the loss and gain of both 9q homologs (9q nullisomy and tetrasomy, respectively, Figure 8D) as well as the frequent loss of one 9q homolog (monosomy of 9q, Figure 8D). The 9q monosomies were not accompanied by an increase in 9q trisomies, which we had expected based on the fixed cell imaging analysis (Figure 7, 8A). Since we randomly sorted whole single G1 cells, that had gone through S phase and mitosis based on EdU incorporation and Hoechst labeling, and did not select for daughter cell pairs, nor for cells with unequal distribution of the 9q locus, we may have accidently under-sorted cells with 9q trisomies. Taken together, the combined live cell imaging, IF and single cell Karyo-seq data suggest that targeting Kin14VIb to endogenous repetitive loci on single chromosomes is a powerful means to mis-segregate chromosomes, providing a tool to investigate the direct consequences of single chromosome mis-segregation events.

**Figure 7:**
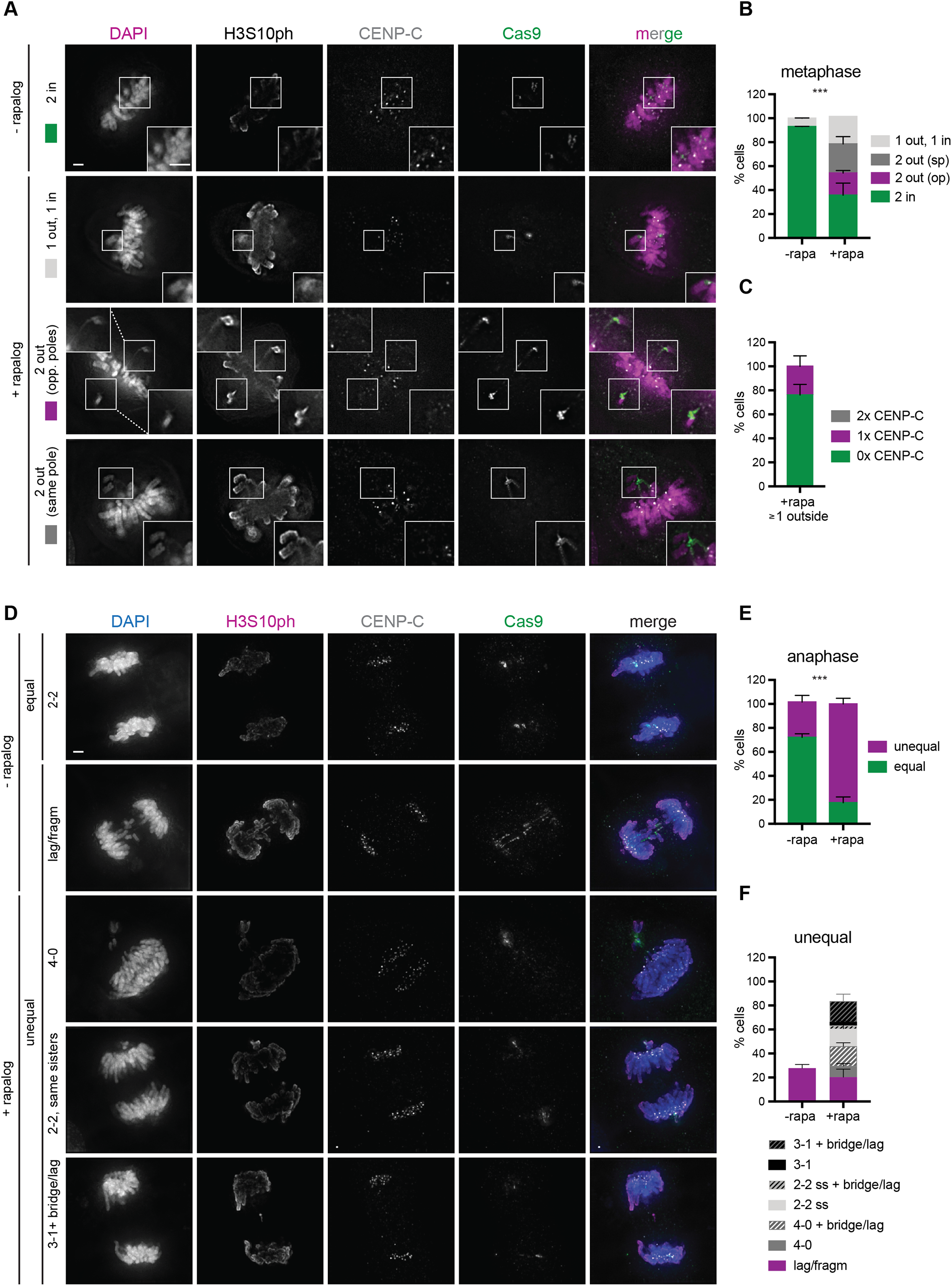
Segregation errors induced after Kin14IVb tethering to Chr9. **A, D**. Representative IF images of RPE1-dCas9-Kin14VIb cells in metaphase (A) or anaphase (D), 3 days after transduction with Chr9-cen sgRNAs and in the presence or absence of rapalog. Insets (A) are magnifications of the white boxed region (scale bar = 2 µm). **B**. Frequency of the observed metaphase localizations of the Chr9-cen loci as shown in (A). *** p<0.001 (Fisher’s exact test, comparing the fraction of cells displaying 2 Chr9-cen loci **in**side vs. at least 1 Chr9-cen locus **out**side the metaphase plate in the absence or presence of rapalog). **C**. Fraction of cells with spindle pole localization of the Chr9-cen loci and CENP-C in close proximity. Experiment was performed in duplicate (mean ± SD, n ≥ 35 per condition, per experiment). **E**. Quantification of anaphase distribution of the Chr9-cen loci *** p<0.001 (Fisher’s exact test, comparing the fraction of cells with equal vs. unequal distribution of Chr9-cen loci in the absence or presence of rapalog). **F**. Frequency of the different types of segregation errors observed in the cases of an unequal distribution of the Chr9-cen loci. Experiment was performed in duplicate (mean ± SD, n ≥ 28 per condition, per experiment). Sp = same pole, op = opposite pole, ss = same sisters.

**Figure 8:**
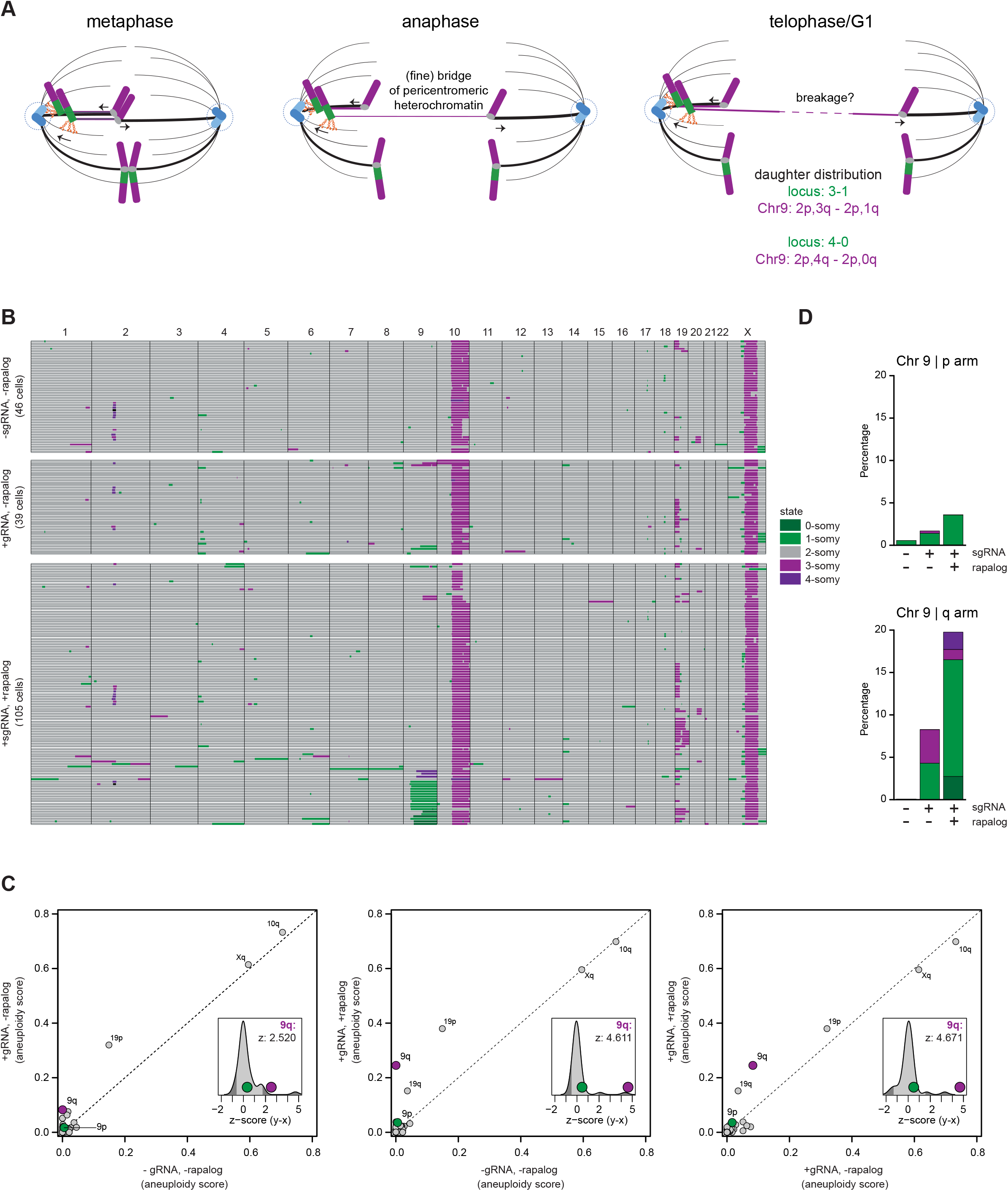
Tethering Kin14VIb to a pericentromeric DNA repeat in Chr9 results in 9q aneuploidies after a single cell division. **A**. Scheme illustrating how Kin14VIb binding to an endogenous pericentromeric repeat in Chr9q may cause Chr9 arm level aneuploidies after a single cell division. For schematic clarity, only the situation where one homologue is at the pole during metaphase is drawn. **B**. Genome-wide copy number heatmaps of RPE1 dCas9-Kin14IVb cells for the indicated conditions. Note that in the -sgRNA condition, Kin14VIb is not expression (-dox), whilst in the +gRNA conditions the kinesin is expressed (+dox). Individual cells are in rows and genome positions in columns. Chromosome boundaries are indicated with black lines and colors are corresponding to the most likely copy number state per bin as determined by AneuFinder. **C**. Plots comparing aneuploidy scores for each chromosomal arm between the various indicated conditions. The aneuploidy scores of the chromosomal arms are deviations from euploid (average absolute difference from 2-somy of the bins that are associated with the arm; see materials and methods). Inserts show density plots of the difference of the scores of the two conditions that are compared (y-axis minus x-axis). These differences are expressed as z-scores (number of standard deviations from the mean). The 95% confidence interval is depicted in light grey. The values of the 9p and 9q arm are indicated with a green and purple dot respectively. Z-score value for 9q is indicated (see also Supplemental Table 2). **D**. Average percentage of of the bases of the p or q arm that are (classified as) 0, 1, 3 or 4-somy. Percentages are depicted as stacked barplots and are determined for all copy numbers other than 2-somy (euploid).

## Discussion

We here demonstrate the feasibility of manipulating the orientation and segregation of specific human chromosomes during mitosis by recruiting the minus-end-directed Kin14VIb of *P. patens* to chromosome-specific repetitive loci. Interestingly, in maize, the so-called abnormal chromosome 10 (Ab10), harbors a cluster of eight genes, known as the Kinesin driver (Kindr) complex, that promotes meiotic drive, i.e. the preferential transmission of Ab10 into egg cells ^53^. The mechanism of Ab10 drive involves the conversion of inert heterochromatic regions called knobs into motile ‘units’ that are actively transported along microtubules toward spindle poles during meiosis I and II ^54^. It was recently demonstrated that Ab10 knob transport is mediated by Kindr encoded KINDR, a minus-end directed motor protein that diverged from a Kinesin 14A ancestor and that specifically interacts with 180 bp knob repeats ^53^. Through TetR and dCas9-mediated tethering of *Pp* Kin14VIb to pericentromeric and subtelomeric heterochromatic repeats, we appear to have mimicked this chromosomal ‘drive’ mechanism in human mitotic cells, forcing unequal transmission of targeted chromosomal arms into daughter cells.

We find that the accumulation of Kin14VIb dimers on a highly repetitive (∼ 200 × 96 mer) integrated TetO locus induces poleward transport of the TetO locus in U-2 OS cells ^33^, which caused the tethered chromosome 1p arm to be pulled towards the spindle pole. We noted, however, that GCN4-induced Kin14VIb tetramers are more effective in chromosome transport than Kin14VIb dimers, when employing dCas9 to tether the motor to endogenous Chr1p36-specific DNA repeats. The lower number of sgRNA binding sites in the endogenous repeats compared to TetR-binding sites in the integrated TetO repeat, will likely result in a smaller number of dCas9 and hence Kin14VIb molecules to accumulate on the endogenous locus, thereby affecting the probability of the dimeric motor to dimerize ^29,55^.

Kin14VIb tethering to either subtelomeric or pericentromeric loci allowed microtubule attachment of the KTs of the targeted chromosome. When tethered to a subtelomeric repeat in Chr1, it even allowed the acquisition of bi-oriented MT attachments by the sister-kinetochores that facilitated sister KT congression. Although we frequently observed poleward orientation of the Kin14VIb bound loci after NEB, it is possible that MT capture of the respective KTs is even faster and was already established before Kin14VIb began transporting the telomere towards the spindle pole. Alternatively, Kin14VIb-mediated telomere transport may have started before the kinetochores acquired MT attachments. The latter scenario would suggest that the movement of the ends of the Chr1 sister p arms towards the spindle pole does not affect the back-to-back orientation of sister KTs that facilitates their bipolar MT capture, although we cannot exclude that the motor-induced p arm pulling may enhance merotelic capture of one of the sister KTs. Even when Kin14VIb was bound to the large pericentromeric repeat of Chr9q, one or two of the nearby sister KTs could attach to MTs derived from the opposing spindle pole causing the pericentromeric heterochromatin to stretch and spatially separate from the kinetochore.

The opposing forces exerted by Kin14IVIb and kinetochore MTs on the same chromosome had striking consequences for the segregation of that chromosome during anaphase. When bound close to the telomere of Chr1, Kin14VIb caused the p arm to stretch heavily when the KT of the sister chromatid was connected to MTs derived from the opposing spindle pole, revealing the plasticity of a human mitotic chromosome under physiological conditions, i.e. in living cells. The increase in Chr1p aneuploidy scores deduced from whole genome sequences of individual U-2 OS nuclei, implies that the stretched 1p arm is eventually broken and gained in or lost from the daughter nuclei. Since mere spindle forces are considered not strong enough to induce DNA double strand breaks ^44,56^, we consider it more likely that, similar to chromatin bridges formed by dicentric chromosomes as a consequence of telomere fusions, the 1p arm gets trapped and damaged in the intercellular cytokinetic bridge during telophase. Future work will show if breakage of the 1p arm involves the action of the endonuclease TREX1 or is mediated by actomyosin forces as described for bridges formed by dicentric chromosomes ^43–45^. When bound to pericentromeric repeats of Chr9q close to the sister-KTs, the opposing pulling forces exerted by kMTs and Kin14VIb appeared to stretch part of the pericentromeric heterochromatin, separating the sister q arms from the p arms and kinetochores during metaphase and anaphase, and resulting in 9q gains and losses in the daughter cells (Figure 8A).

In conclusion, we have developed a molecular motor-based approach that allows both the visualization and manipulation of specific chromosomes during mitosis and by which we can induce chromosome-specific segmental aneuploidies in daughter cells after one round of cell division. In an accompanying manuscript, Tovini *et al*. employed dCas9-CENP-T fusion proteins to build an ectopic kinetochore on targeted chromosomes, a strategy that also increased the frequency of chromosome-specific CIN and aneuploidies. When for instance combined with microscopy-based single-cell isolation methods and subsequent sequencing, such as described for Look-Seq ^57^, Live-Seq ^58^, or Photostick ^59^, our methods open up the possibility to investigate the immediate cellular responses to specific (arm level) aneuploidies in different cell types; an important step towards understanding how recurrent aneuploidy patterns arise in different cancer types. Moreover, by taking advantage of the poleward pulling forces exerted by Kin14VIb on the targeted chromosome, our method can also be applied to study biophysical properties of mitotic human chromosomes in living cells.

## Acknowledgements

We thank the Oncode Single-cell (epi) genome sequencing facility for scKaryo-Seq, Dr. M. Hadders for help with figures 5 and 6, all scientists that shared their plasmids via Addgene, and Sarah McClelland for helpful discussions and sharing unpublished data. This work is part of Oncode Institute, which is partly financed by the Dutch Cancer Society. This project was co-funded by a Dutch Cancer Society grant (2018-RUG-11457) to F.F. Further support came from the European Research Council (ERC Consolidator Grant 819219 to L.C.K.) and the Centre for Living Technologies, a part of the Alliance TU/e, WUR, UU, UMC Utrecht (www.ewuu.nl).

## Materials and Methods Cells and cell culture

HEK293T cells were cultured in DMEM (Sigma-Aldrich) medium. U-2 OS TetO (Janicki et al., 2004) and all RPE1-TERT cell lines were cultured in DMEM-F12 medium (Sigma-Aldrich). Medium was supplemented with 10% fetal bovine serum (Bodinco B.V.), 1 mM ultraglutamine (Lonza), and 0.1 mg/ml penicillin/streptomycin (Sigma-Aldrich). Cells were cultured at 37°C and 5% CO_2_ in a humidified incubator.

## Plasmids

The lentiviral vector F9-rTetR-EGFP-IRES-PuroR was generated by Gibson cloning from F9-TetR-EGFP-IRES-PuroR (gift from Huimin Zhao, Addgene plasmid # 117049). TetR was replaced by reverse (r) TetR obtained by PCR from AAVS1 TRE3G-GFP (gift from Su-Chun Zhang, Addgene plasmid # 52343). *Pp*Kin14VIb cDNA was obtained by PCR from pSIN-TRE-rtTA-IRES-Puro-FRB-mCherry-GCN4-*Pp*Kin14VIb and cloned into F9-rTetR-EGFP-IRES-PuroR to generate F9-rTetR-EGFP-*Pp* Kin14VIb-IRES-PuroR. pSIN-TRE-rtTA-IRES-PuroR-FRB-mCherry-GCN4-*Pp*Kin14VIb(861-1321) was constructed from pSIN-TRE-rtTA-IRES-Puro (kindly provided by Benjamin Bouchet, Utrecht University, The Netherlands), and encodes tetramerized GCN4-*Pp* Kin14VIb (aa 861–1321), derived from the plasmid *Pp*Kin14VIb–GCN4, a gift from Gohta Goshima (Nagoya University, Nagoya, Japan), and fused N-terminally to the rapalog-sensitive heterodimerization module FRB and mCherry. pSIN-TRE-rtTA-IRES-PuroR-FRB-mCherry-GCN4-*Pp*Kin14VIb(861-1321) is a self-inactivating lentiviral vector enabling doxycycline-sensitive expression of the minus-end directed kinesin FRB-mCherry-GCN4-*Pp*Kin14VIb(861-1321), carrying a puromycin resistance cassette and the reverse tetracycline-controlled transactivator, rtTA.

To generate the lentiviral vector pCuO-rTetR-EGFP, rTetR-EGFP was amplified by PCR from F9-rTetR-EGFP-IRES-PuroR and cloned into pCDH-CuO-MCS (SystemBio, cat. no. QM500A-1) by Gibson assembly. Human codon-optimized *Pp*Kin14VIb (Integrated DNA Technologies) was subsequently cloned into pCuO-rTetR-EGFP to generate pCuO-rTetR-EGFP-*Pp*Kin14VIb.

The lentiviral vector pHAGE-UbC-dCas9-GFP-3xFKBP was generated from pHAGE-TO-dCas9-3xGFP, a gift from Thoru Pederson (Addgene plasmid # 64107). By Gibson cloning, the final 2xGFP were replaced by 3xFKBP, obtained by PCR from pcDNA5 FKBP-GFP (a gift from Geert Kops, Hubrecht Institute, the Netherlands). The Chr1-telo (sequence: GATGCTCACCT) and Chr9-cen (sequence: TGGAATGGAATGGAATGGAA) sgRNAs were selected based on the principles described in ^50^, and cloned into lentiviral vector pLH-spsgRNA2 (gift from Thoru Pederson, Addgene plasmid # 64114). The Chr1-telo sgRNA was used by us in Dumont *et al*. ^8^.

For CRISPR/Cas9 knockout of the puromycin N-acetyl-transferase (PAC) gene in RPE1 cells, the sgRNA GGCGGGGTAGTCGGCGAACG was cloned into the expression vector pAceBac1-U6-CBA-Cas9-2A-EGFP, generated as previously described ^60^.

## Lentivirus production and transduction

For lentivirus production, HEK293T were seeded in 10 cm dishes at 10% confluency 24 hours prior to transfection. Cells were co-transfected with lentiviral vectors plus the lentiviral packaging and envelope plasmids: psPAX2 (Addgene plasmid #12260) and psMD2.G (Addgene plasmid #12259). Lentivirus containing FRB-mCherry-GCN4-Kin14VIb and rTetR-GFP-Kin14VIb were harvested by filtering the cell supernatant through a 40 µm filter at 48 and 72 hours post-transfection, and were incubated in a 5x concentrating solution consisting of 400g/L PEG 6000 and 0,41M NaCl (pH7.2) at 4°C overnight. The supernatant and concentrating solution mixture was centrifuged at 4500 rpm for at least 30 minutes, resuspended in 100 ul ice-cold PBS, and stored at -80°C until transduction. Lentivirus containing dCas9-GFP-3xFKBP, rTetR-GFP, chr1-telo sgRNA and chr9-cen sgRNA were harvested 48 hours post-transfection as described above and stored at -80°C until transduction. For lentiviral transduction, cells were seeded at 30%–40% confluency and incubated 16 hours with either 100, 500 or 1000 µl lentivirus containing FRB-mCherry-GCN4-Kin14VIb or rTetR-GFP-Kin14VIb, chr1-telo or chr9-cen sgRNAs, and dCas9-GFP-3xFKBP or rTetR-GFP, respectively, in the presence of 6 µg/ml polybrene.

## Cell line generation

To generate RPE1 cell lines stably expressing dCas9-GFP-3xFKBP, RPE1 cells were transduced with lentivirus containing dCas9-GFP-3xFKBP, and after a week were sorted for low GFP expression by flow cytometry. Subsequently, RPE1 cells stably expressing dCas9-GFP-3xFKBP were transduced with lentivirus containing FRB-mCherry-GCN4-Kin14VIb. A population of high mCherry-Kin14VIb-expressing cells was obtained by flow cytometry, expanded in culture, and plated as single cells into 96-well plates containing 100 puromycin-sensitive feeder RPE1 cells. Puromycin-sensitive RPE1 cells were generated by CRISPR-Cas9 mediated mutation of the puromycin N-acetyl-transferase (PAC) gene. Seven days after plating, feeder cells were eliminated by puromycin selection (1µg/ml, Sigma-Aldrich). Expanded clones were transduced with lentivirus containing chr1-telo sgRNA and treated for 16 hours with 20 µM of the Eg5 inhibitor S-trityl-L-Cysteine (STLC, Tocris Biosciences) to generate a population of mitotic cells with monopolar spindles. Cells were then treated with 500 nM rapalog (AP21967/AC heterodimerizer, Clontech) to induce FKBP12-FRB dimerization, and were followed by live cell imaging. The clone with the largest fraction of cells showing polar localization of chr1-telo GFP foci after rapalog addition was selected for further experiments. This cell line is referred to as RPE1 dCas9-Kin14VIb.

## Live cell imaging

U-2 OS TetO cells stably expressing H2B-mCherry were transduced with lentivirus containing rTetR-GFP or rTetR-GFP-Kin14VIb. 9 days post-transduction, cells were seeded in an optical-quality plastic 8-well slide (IBIDI, cat. no. 80826) for live cell imaging. The next day, media was replaced by FluoroBrite DMEM (Gibco) supplemented with 10% fetal bovine serum (Bodinco B.V.), 1mM ultraglutamine (Lonza), and 0.1 mg/ml penicillin/streptomycin (Sigma-Aldrich). Cells were treated with 1 µg/ml doxycycline to induce TetO-rTetR binding and immediately imaged live. Images were acquired every 5 minutes on a Zeiss AIM System - Cell Observer microscope equipped with an AxioImager Z1 stand, a Hamamatsu ORCA-flash 4.0 camera, and a Colibri 7 LED module, using a 40×/1.4 oil PLAN Apochromat lens. All movies were subsequently processed and analyzed using ZEN software (Zeiss).

RPE1 dCas9-Kin14VIb cells were transduced with lentivirus containing chr1-telo or chr9-cen sgRNAs. 24 hours after transduction, cells were plated for live cell imaging in an optical-quality plastic 8-well slide (IBIDI, cat. no. 80826). The next day, cells were incubated with 200 nM SiR-DNA (Spirochrome) and 1µg/ml doxycycline to visualize DNA and induce FRB-mCherry-Kin14VIb expression, respectively. After 8 hours, cells were treated with 500 nM rapalog (Clontech) to induce FRB-FKBP dimerization, and images were acquired every 5 minutes on a Zeiss AIM System - Cell Observer microscope as described above.

## Chromosome spreads

For chromosome spread preparations, cells were synchronized in mitosis by a 16 hour incubation in 20 uM STLC (Tocris Biosciences). Following a 10 minutes treatment with 0.83 µM nocodazole (Sigma-Aldrich), mitotic cells were harvested by shake-off and incubated 10 minutes in 75 mM KCl. Subsequently, cells were spun onto glass coverslips in a Cytospin centrifuge at 1500 rpm for 5 minutes. Samples were fixed and stained as described below.

## Cell synchronization and immunofluorescence

U-2 OS TetO cells were transduced with lentivirus containing rTetR-GFP or rTetR-GFP-Kin14VIb. 7 days post-transduction, cells were plated on coverslips for immunofluorescence and incubated overnight in 7 uM RO-3306 (Sigma-Aldrich) to induce a G2 arrest. The next day, cells were washed 3x with warm media to release from RO-3306 and incubated 1 hour in medium containing 1µg/ml doxycycline to induce TetO-rTetR binding, prior to fixation. Alternatively, cells were synchronized in metaphase by a 20 min RO-3306 release in the presence of 1µg/ml doxycycline, followed by a 45 min block in 5 uM MG-132 (Calbiochem). RPE1 dCas9-Kin14VIb cells were transduced with lentivirus containing Chr1-telo or Chr9-cen sgRNAs and 48 hours later plated in an optical-quality plastic 8-well slide (IBIDI, cat. no. 80826) for immunofluorescence. To induce Kin14VIb expression, RPE1 were treated with 1µg/ml doxycycline for at least 8 hours.

All samples were fixed in 4% PFA for 10 minutes and permeabilized with ice-cold methanol for at least 10 minutes, unless otherwise stated. For Mad1 IF, U-2 OS TetO cells were plated onto poly-D-Lysine-coated coverslips, synchronized in metaphase as described above, and fixed and permeabilized simultaneously in 4% PFA diluted in PHEM-Triton buffer (60mM HEPES, 20mM Pipes, 10mM EGTA, 2mM MgCl_2_, 0.2% Triton, pH 6.9). Fixed samples were blocked in 3% BSA solubilized in PBS containing 0.1% Tween 20, incubated with primary antibodies for 2 hours, washed, and incubated with secondary antibodies and 1µg/ml DAPI (Sigma-Aldrich) for 1 hour. For γH2AX IF, samples were fixed in 4% PFA for 15 minutes and incubated in PBS-Triton X-100 (0.5%) for 10 minutes to retrieve antigens. Blocking was performed in 2.5% BSA solubilized in PBS-Tween 20 (0.05%) for 15 minutes, and samples were incubated with primary antibodies at 4°C overnight. Secondary antibody incubation was performed for 2 hours at RT. The following primary antibodies were used: GFP booster (Chromotek, gba488), mouse anti-yH2AX (Millipore 05-636), mouse anti-H3S10ph (Millipore, 05-806), rabbit anti-H3S10ph (Millipore, 06-570), guinea pig anti-CENP-C (MBL, PD-030), mouse anti-Mad1 (Millipore, MABE867), rabbit anti-pericentrin (Abcam, ab4448), mouse anti-Cas9 (Diagenode, C15200203), rabbit anti-RFP (Rockland, ROCK600-401-379). Coverslips were mounted with Prolong Diamond (Invitrogen), and all samples were imaged with an inverted 100x oil objective on an Olympus IX71 microscope connected to a Deltavision imaging system and a CoolSnap HQ camera (Photometrics).

## FISH

U-2 OS TetO cells were transduced with lentiviruses containing rTetR-GFP or rTetR-GFP-Kin14VIb, replated at 60% confluency 48 hours post-transduction and incubated in 7 µM RO-3306 overnight. The next day, cells were washed 3x with warm media and incubated 1 hour in medium containing 1ug/ml doxycycline to induce TetO-rTetR binding, prior to fixation. Cells were fixed in MetOH:acetic acid (3:1) for 10 minutes at -20°C, and FISH using Chromosome 1 Classical Satellite Probe (LPE001R, Cytocell) was performed according to manufacturer’s instructions.

## Single cell karyotype sequencing

U-2 OS TetO cells were transduced with lentiviruses containing rTetR-GFP or rTetR-GFP-Kin14VIb. After 48 hours, cells were replated at 60% confluency and incubated in 7 µM RO-3306 overnight. The next day, cells were washed 3x with warm media and incubated in medium containing 1µg/ml doxycycline to induce TetO-rTetR binding. 4 hours after RO-3306 release, single nuclei were extracted by incubating cells on ice for at least 15 minutes in nuclei suspension buffer (NSB) containing 0.1 M Tris-HCl pH 7.5, 0.154 M NaCl, 1 mM CaCl_2_, 0.5 mM MgCl_2_, 0.2 % BSA, 0.1% NP40, and 1µg/ml Hoechst 34580 (Sigma-Aldrich).

RPE1 dCas9-Kin14VIb cells were transduced with lentivirus containing chr9-cen sgRNAs. After 16 hours, cells were treated with 1µg/ml doxycycline to induce FRB-mCherry-GCN4-Kin14VIb expression. 24 hours post-transduction, cells were replated into a 6-well plate at a density of 170.000 cells/ml, and incubated with 5 µM EdU (Thermofisher) +/-500 nM Rapalog (Clontech) for 17 hours. Cells were then fixed using 70% ice-cold ethanol, washed with 1x Saponin-based permeabilizing and washing reagent (Thermofisher), and incubated for 30 minutes with the Click-iT reaction cocktail according to the manufacturer’s protocol (Thermofisher). Cells were washed with 1x Saponin-based permeabilizing and washing reagent, and DNA was stained using 1µg/ml DAPI (Sigma-Aldrich).

Single nuclei or single cells were filtered through 40 μm strainers, and sorted into 384-well plates containing 5 ul mineral oil (Sigma-Aldrich). Samples were further processed as described previously (Bolhaquero et al., 2019). Libraries of U-2 OS TetO and RPE1 cells were sequenced on an NovaSeq6000 S1 2×100bp (80M reads/384 single cells) and NovaSeq6000 S1 1×100bp (67M reads/384 single cells) sequencer, respectively. The fastq files were mapped to GRCH38 using the Burrows–Wheeler aligner. The aligned read data (bam files) were analyzed with a copy number calling algorithm called AneuFinder (version 1.14.0; ^61^. Following GC correction and blacklisting of artefact-prone regions (extreme low or high coverage in control samples), libraries were analyzed using the dnacopy and edivisive copy number calling algorithms with variable width bins (average binsize = 1 Mb; step size = 500 kb). Breakpoints were refined as well (refine.breakpoints = TRUE). The samples were analyzed with an euploid reference ^62^. Results were afterwards curated by requiring a minimum concordance of 90 % between the results of the two algorithms. Libraries with on average less than 10 reads per bin and per chromosome copy (∼55,000 reads for a diploid genome) were discarded. The dnacopy results are presented. The copy numbers of the U-2 OS TetO samples were expressed relative to the median (consensus) copy number profile of the non-transduced U-2 OS TetO cells (“non-transduced”) in order to account for the non-euploid nature of this cell line (see Figure S3A). This median copy number was determined for each bin (median across libraries). Aneuploid scores were first calculated for each bin by calculating the average absolute difference from this median copy number profile (average across libraries). Scores for chromosomal arms were subsequently calculated by averaging the scores of the bins that were associated with each arm (weighted average; bins have variable width). The aneuploidy scores of the RPE1 samples were calculated in the same way except that the scores were calculated relative to an euploid reference (absolute difference from 2-somy).

## Statistics

Where indicated, the mean and standard deviation (SD) are shown. Statistical significance was calculated with a χ^2^ test, Fischer exact test, or non-parametric Student’s t test (Mann-Whitney test) using Prism 8 software.

## Figure legends

**Figure S1 (related to Figure 2): Mad1 kinetochore localization in U-2 OS TetO cells**

**A**. IF for GFP, Mad1 and CENP-C of U-2 OS-TetO cells in metaphase expressing rTetR-GFP (ctrl) or rTetR-GFP-Kin14VIb (Kin14VIb). Representative images of the different orientations of the TetO chromosome or locus and the sister KTs are shown. DNA visualized by DAPI. Scale bar = 2 µm. **B**. Schematics of amphitelic, merotelic, and syntelic kinetochore-microtubule attachment states. Only the amphitelic attachment state results in error-free segregation of the sister chromatids.

**Figure S2 (related to Figure 3): Stretching of the Kin14VIb-bound TetO chromosome during anaphase and telophase**.

**A**. Representative images of a U-2 OS-TetO cell expressing TetR-GFP-Kin14VIb displaying a 2-0 distribution of the TetO loci and two stretched arms along the mid-axis (2 stretched arms, figure 3B). Scale bar = 5 µm. **B**. Scheme illustrating potential metaphase positioning and anaphase segregation of the TetO chromosome as shown in (A). While Kin14VIb pulls the p arm telomeres of the TetO chromosome towards one pole, its sister KTs are syntelically attached to MTs coming from the opposite spindle pole, thereby causing the two sister arms to stretch across the two spindle poles. **C**. IF for CENP-C, GFP, H3S10ph of U-2 OS TetO cells in anaphase expressing rTetR-GFP-Kin14VIb. DNA visualized with DAPI. Examples of heavily stretched Chr1p arms caused by transport of the telomere towards one spindle pole by Kin14VIb and attachment of the KT by MTs coming from the opposite pole. White arrowhead indicates the kinetochore (CENP-C) of the stretched chromatid. **D**. IF for Aurora B, GFP, H3S10ph of U-2 OS TetO cells in anaphase and telophase expressing rTetR-GFP-Kin14VIb. The stretched chromatid arm (visible with H3S10ph) gets trapped into the cytokinetic furrow and midbody, highlighted by the respective midzone and midbody localization of Aurora B. Scale bar = 5 µm.

**Figure S3 (related to Figure 4). Single nuclei karyotypes of U-2 OS TetO cells**

**A**. Genome-wide copy number heatmap of the non-transduced U-2 OS TetO cells (non-transduced). Individual cells are in rows and genome positions in columns. Chromosome boundaries are indicated with black lines and colors are corresponding to the most likely copy number state per bin as determined by AneuFinder. The median copy number was determined for each bin (median across libraries) to produce a median (consensus) karyotype (colored line below heatmap). **B**. The latter was used as reference to generate heatmaps showing the copy number alterations relative to the U-2 OS TetO median karyotype for non-transduced (upper), rTetR-GFP-(middle), or rTetR-GFP-Kin14VIb-transduced (lower) U-2 OS TetO cells.

**Figure S4 (related to Figure 3, 4). γH2AX is not detected on the TetO locus nor on the H3S10ph+ bridges in U-2 OS TetO cells expressing rTetR-GFP-Kin14VIb**

**A**. Representative IF images of U-2 OS TetO cells in anaphase/telophase expressing rTetR-GFP (ctrl) or rTetR-GFP-Kin14VIb (Kin14VIb) and stained with the indicated antibodies. γH2AX was used as a marker for DNA damage. Magnifications of the white boxed regions are shown in the corners of the image images. Solid lined boxes represent the GFP (TetO) foci, the dotted lined boxes the chromatin bridges. Scale bars = 5 µm. **B**. Quantification of gH2AX co-localization with the segregating TetO locus in U-2 OS cells expressing rTetR-GFP or rTetR-GFP-Kin14VIb. **C**. Frequency of γH2AX detection on chromatin bridges (incl. stretched arms) and lagging chromosomes during anaphase/telophase. N = the number of cells.

