## Supplementary figures and images for "A motor-based approach to induce chromosome-specific mis-segregations in human cells"

### Supplemental Figures

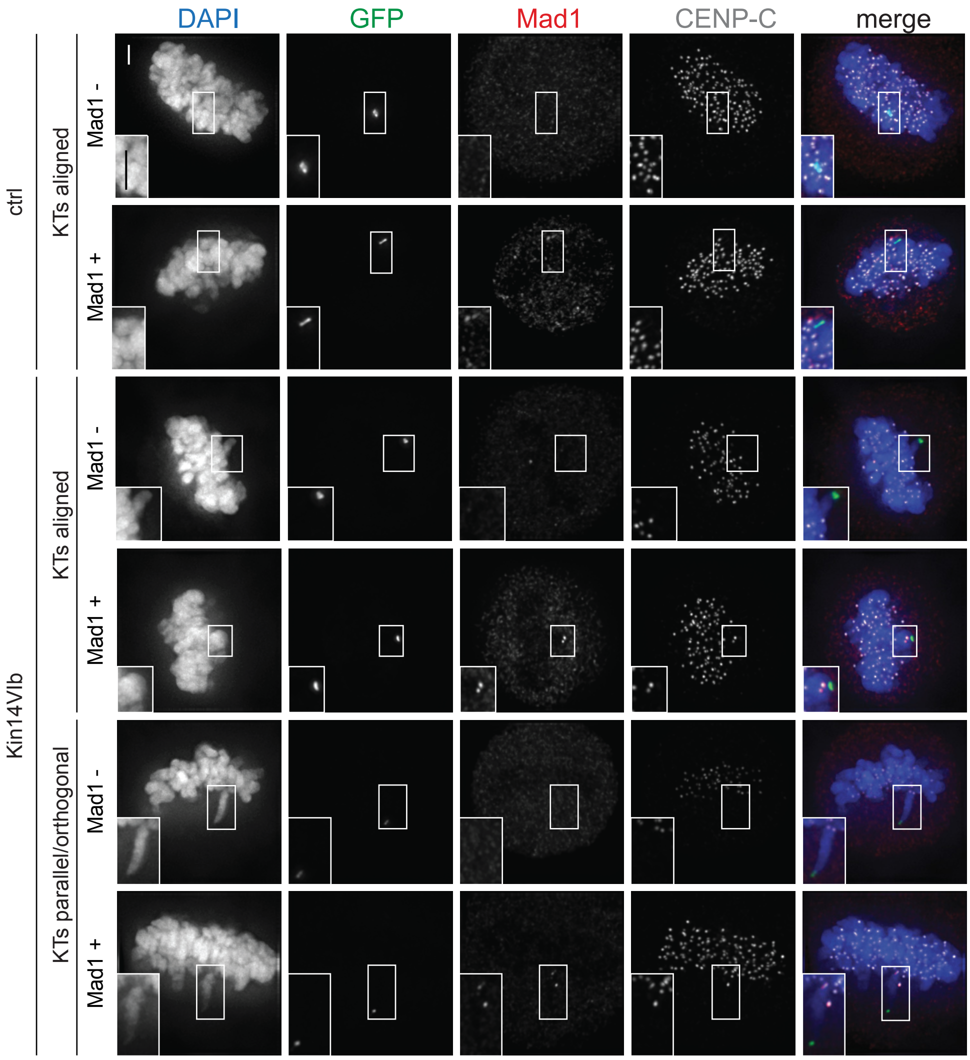

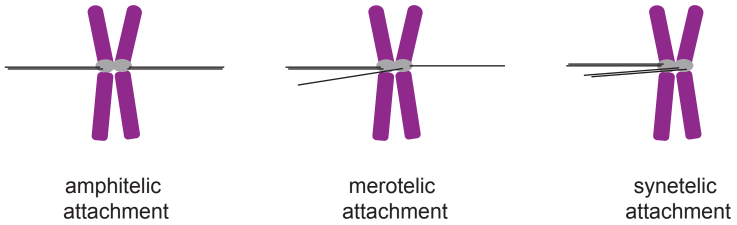

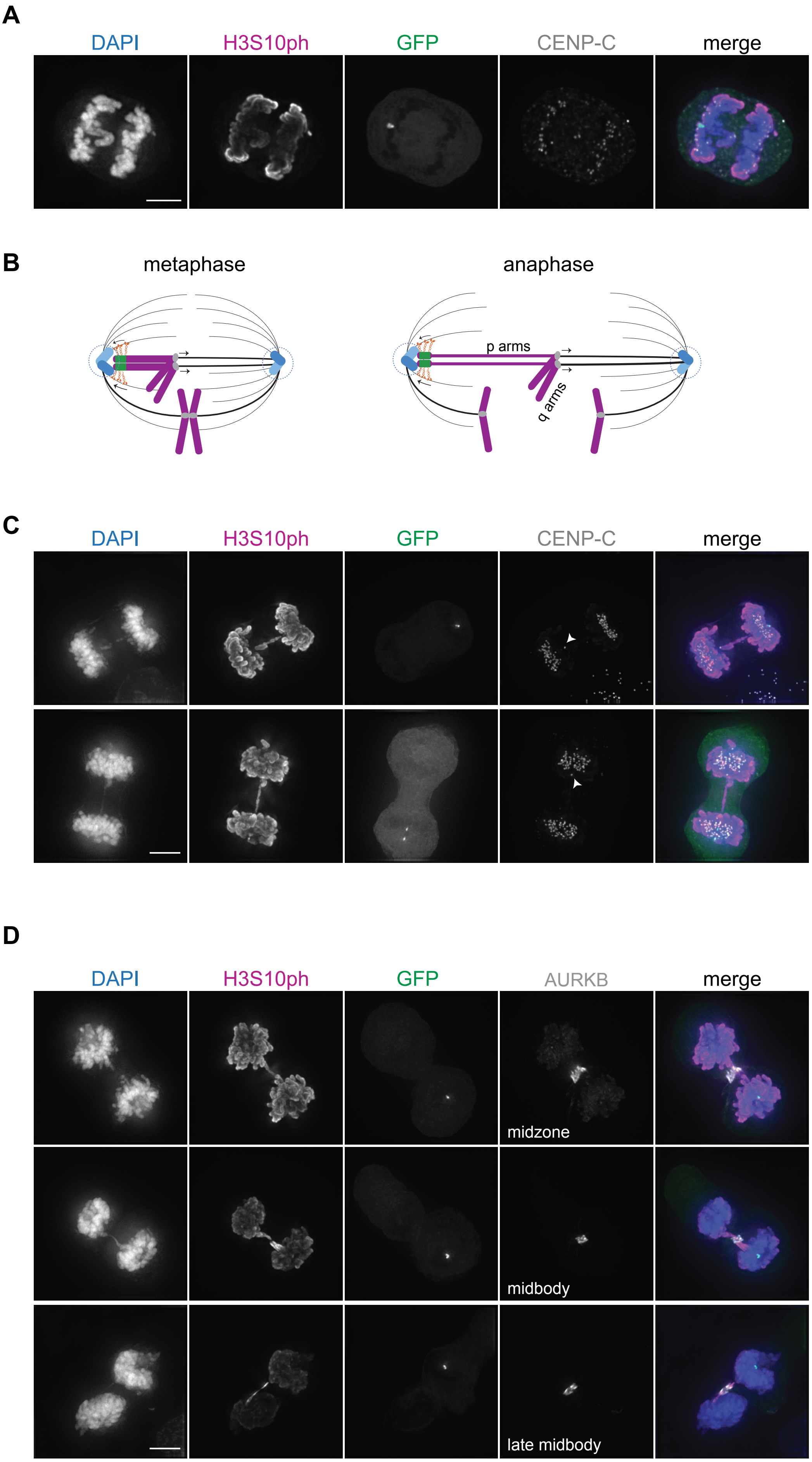

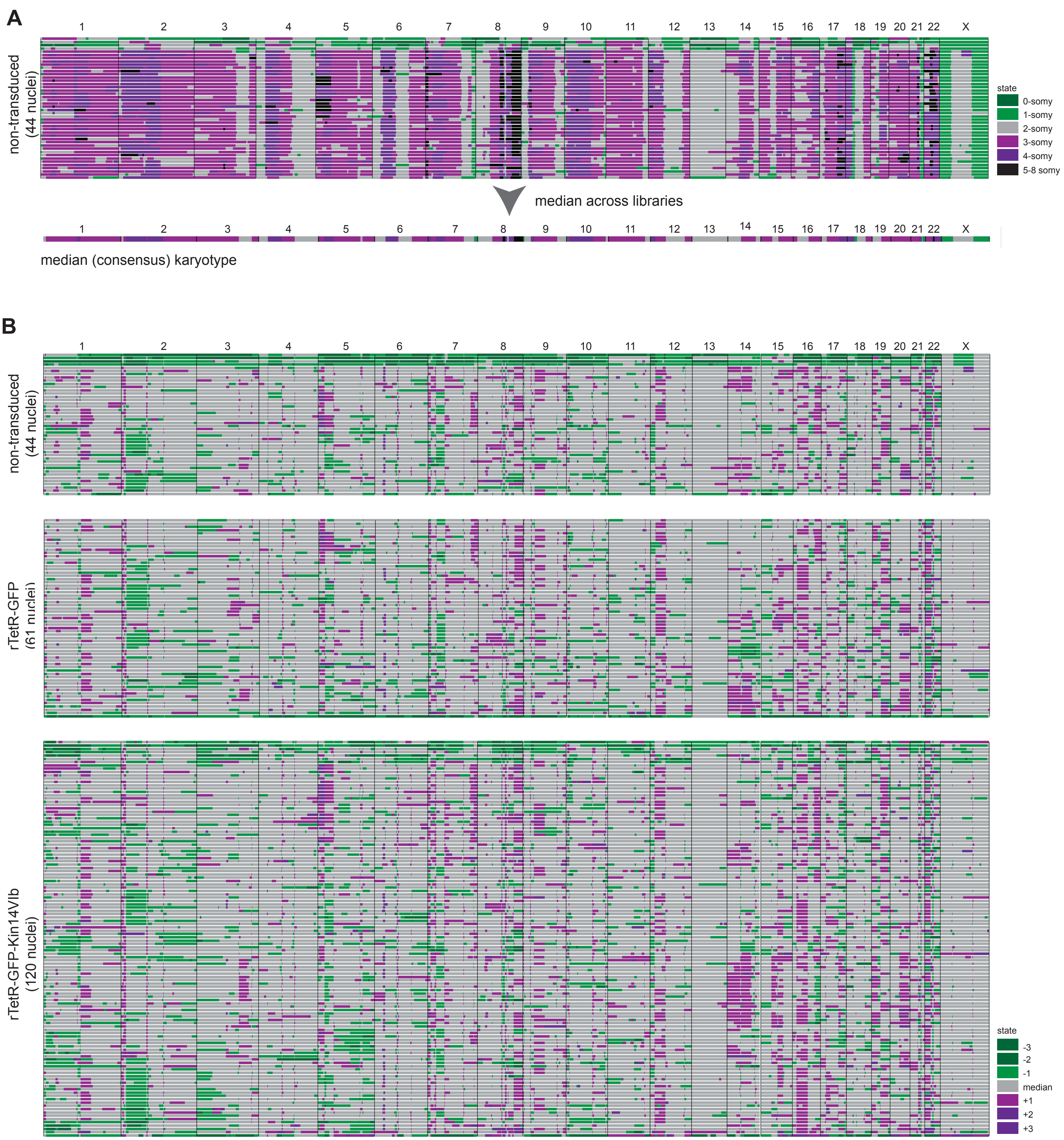

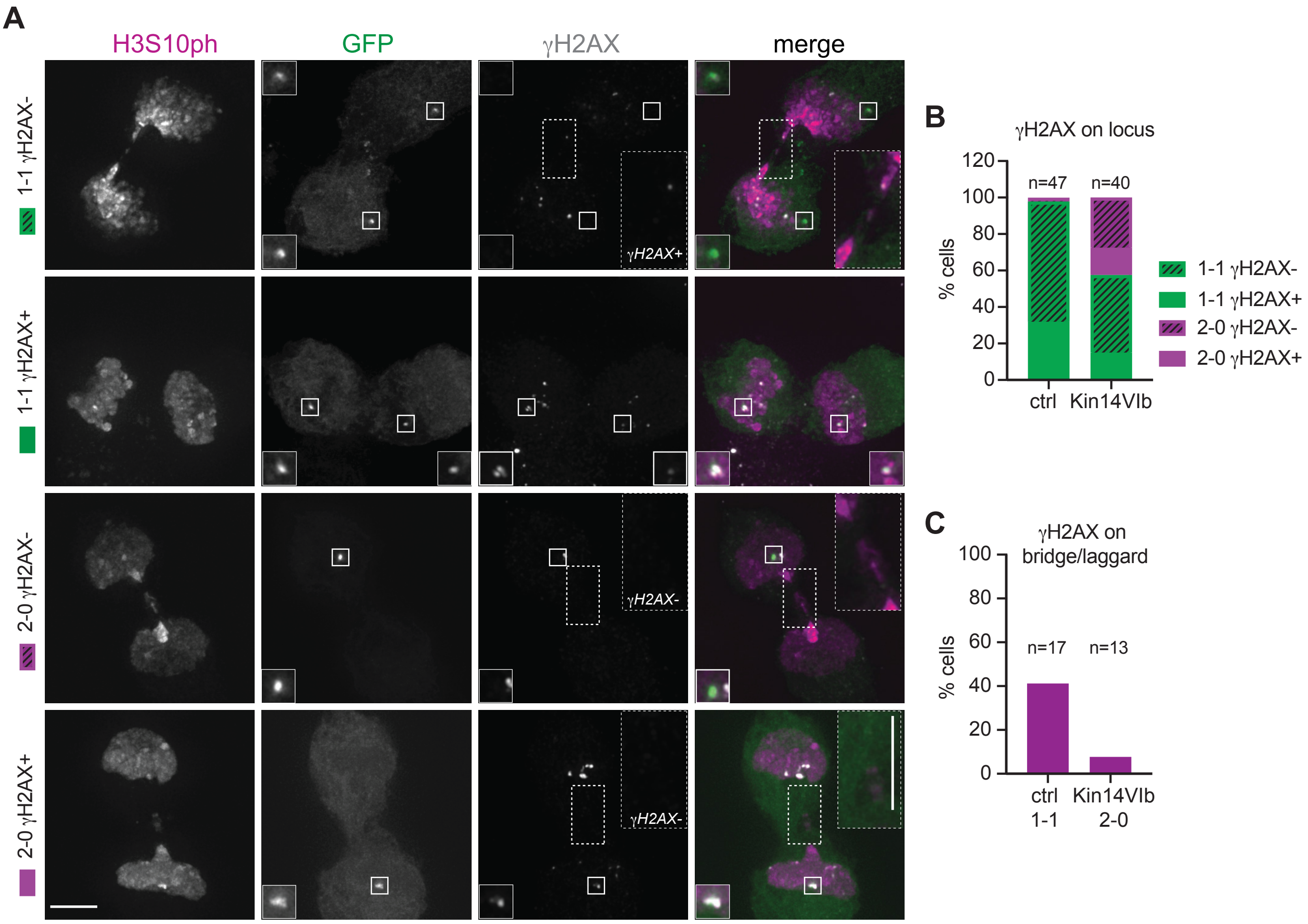
